# Differential genetic variation underlying Ammonium and Nitrate responses in *Arabidopsis thaliana*

**DOI:** 10.1101/2022.04.27.489730

**Authors:** Ella Katz, Anna Knapp, Mariele Lensink, Jordan Stefani, Jia-Jie Li, Emily Shane, Kaelyn Tuermer-Lee, Arnold J. Bloom, Daniel J. Kliebenstein

**Author notes:** Author contributions: EK, DJK and AJB designed the research and wrote the paper, EK, AK, ML, JS, JJL, ES, and KTL performed the research and analyzed the data. The author(s) responsible for distribution of materials integral to the findings presented in this article in accordance with the policy described in the Instructions for Authors (https://academic.oup.com/plcell/pages/General-Instructions) are: Ella Katz and Daniel J. Kliebenstein.

## Abstract

Nitrogen is an essential element required for plant growth and productivity. Understanding the mechanisms and natural genetic variation underlying nitrogen use in plants will facilitate engineering plant nitrogen use to maximize crop productivity while minimizing environmental costs. To understand the scope of natural variation that may influence nitrogen use, we grew 1135 *Arabidopsis thaliana* natural genotypes on two nitrogen sources, nitrate and ammonium, and measured both developmental and defense metabolite traits. By using different environments and focused on multiple traits, we identified a wide array of different nitrogen responses. These responses are associated with a large number of genes, most of them not previously associated with nitrogen responses. Only a small portion of these genes appear to be shared between environments or traits while most of the detected genes are predominantly specific to a developmental or defense trait under a specific nitrogen source. Finally, by using a large population we were able to identify unique nitrogen responses, like preferring ammonium or nitrate, that appear to be generated by combinations of loci rather than a few large effect loci. This suggests that it may be possible to obtain novel phenotypes in complex nitrogen responses by manipulating sets of genes with small effects rather than solely focusing on large effect single gene manipulations.

**One Sentence Summary:** Using a large collection of natural genotypes, and studying both developmental and metabolic responses, we found a large number of genes that are involved in the plants nitrogen response.

## Introduction

Nitrogen is an essential element required for plant growth and productivity. Most plants acquire nitrogen from the rhizosphere either in the form of nitrate (NO_3_^−^) or ammonium (NH_4_^+^) via root absorption (Bloom, 2015). An individual plant’s optimal nitrogen source and concentration range depends on multiple parameters, including plant species, genetics, developmental stage, and the surrounding environment (Britto and Kronzucker, 2002; Fuertes-Mendizábal et al., 2013; Waidmann et al., 2020). Excess application of nitrogen will affect not only the plants growth, but might lead to environmental degradation like groundwater contamination, eutrophication of freshwater, or soil salinization and might accelerate climate change (Zhang et al., 2015). A better understanding of the plant nitrogen acquisition (uptake, assimilation, and regulation thereof) will facilitate adjustments of nitrogen form and concentration to maximize crop productivity yet minimize environmental damage.

Because nitrate is the predominant nitrogen form in temperate agricultural soils, the preponderance of genetic and physiological studies on plant nitrogen relations focuses on nitrate. This focus is based on most plants preferring nitrate as a nitrogen source due to the tendency of ammonium to become toxic if it accumulates in the plant’s cells (Esteban et al., 2016; Liu and von Wirén, 2017; Li et al., 2014; Marino and Moran, 2019). Another potential reason for plant dependence on nitrate is that plants may compete with microorganisms for soil ammonium that microorganisms use as both a nitrogen and energy source (Matson et al., 1998). Nonetheless, preference for nitrate is not ubiquitous and can shift due to genetic variation between and within plants and is sensitive to environmental variation, such as changes in CO_2_ levels (Britto and Kronzucker 2002; Marino and Moran 2019; Menz et al. 2018; Bloom et al., 2012). This suggests that the balance between nitrate vs ammonium usage is not fixed within a plant species, and identifying genetic mechanisms that influence this balance would allow reprogramming plants, by breeding or synthetic biology, to achieve a better match with available nitrogen sources.

To optimize nitrogen acquisition, plants modify root growth and branching, rapidly responding to changes in nitrogen source or concentration. Any shift in the nitrogen condition will typically lead to alterations in primary and lateral root growth and development that maximize the ability to search for more optimal soil nitrogen conditions (Waidmann et al. 2020; Gruber et al. 2013; Gifford et al. 2013; Tian et al. 2009; Rogato et al. 2010; Liu et al. 2013; Drew 1975; Lima et al. 2010; Zhang and Forde 1998; Zhang et al. 1999; Fuertes-Mendizábal et al., 2013; Epstein and Bloom, 2005). Although the mechanisms behind these responses are becoming understood, how many of them are species specific and how many are shared among plants are still unknown.

Shifts in nitrogen source and concentration, in addition to altering root development, can lead to a global reprogramming of the plant’s nutrient uptake, cell metabolic homeostasis, signaling and hormone pathways, defense responses, and responses to elevated CO_2_ (Marino and Moran, 2019; Marino et al., 2016; Bloom et al., 2010). One defense response that is highly dependent on nitrogen availability is production of glucosinolates (GSLs), a class of specialized defense metabolites in *Arabidopsis thaliana* and other plants in the order Brassicales. The breakdown products of GSLs are toxic to herbivores and pathogens and thus have a central role in plant defense against attackers (Beekwilder et al., 2008); (Hansen et al., 2008). GSLs display extensive variation across *Arabidopsis* genotypes, and their composition and accumulation depend on genetics, developmental stage, and external cues (Bakker et al., 2008; Benderoth et al., 2006; Brachi et al., 2015; Chan et al., 2010; Daxenbichler et al., 1991; Halkier and Gershenzon, 2006; Kerwin et al., 2015; Kliebenstein et al. 2001; Kliebenstein et al., 2001b; Kliebenstein et al., 2001a; Rodman, 1980; Sønderby et al., 2010; Wright et al., 2002; Katz et al., 2021). Because GSLs are derived from amino acids and contain nitrogen as part of their basic structure, GSL accumulation is positively correlated with nitrogen supply (Marino et al., 2016; Yan and Chen, 2007; Omirou et al., 2009; He et al., 2014). Furthermore, one plant strategy for dealing with excess ammonium and avoiding toxicity is to accumulate and store more GSLs (Marino and Moran, 2019; La, 2013; Marino et al., 2016; Coleto et al., 2017).

Interestingly, nitrogen and GSLs are bidirectionally coordinated with GSLs influencing nitrogen signaling and/or responses. A metabolic genome-wide association study measuring natural variation in amino acid content within *Arabidopsis* seeds found that two key GSL biosynthetic loci were causally associated with the level of free glutamine in the seed (Slaten et al., 2020). One possible explanation for this GSL to nitrogen connection comes from the observation that specific GSLs can function as growth and development regulators through interactions with different mechanisms including the TOR complex that integrates nitrogen availability into complex signaling networks (Katz et al., 2020; Katz et al., 2015; Malinovsky et al., 2017; Salehin et al., 2019). Similarly MYB29, a GSL transcription factor, can influence root plasticity in response to changes in nitrate levels (Gaudinier et al., 2018). While these studies highlight the involvement of GSLs in nitrogen responses, the mechanisms behind the interplay between nitrogen and defense metabolism are largely unknown.

To develop a deeper understanding of natural variation in *A. thaliana* responses to different nitrogen sources and concentrations, we phenotyped 1135 *Arabidopsis thaliana* natural genotypes (Kawakatsu et al., 2016; The 1001 Genomes Consortium, 2016). While natural variation in nitrogen responses in *Arabidopsis* has received some study, the majority of the work on nitrogen responses in *Arabidopsis* have focused on the reference Columbia-0 (Col-0) genotpye (Menz et al., 2018), and the effectiveness and necessity of using populations with natural variation when querying for disparate responses, i.e. genotypes that prefer ammonium, is under debate (Pigliucci et al., 2006; Waddington, 1953; Pigliucci and Murren, 2003; Schlichting and Pigliucci, 1993). One line of thinking suggests that Col-0 mutants can mimic the entire range of phenotypes available within the species natural variation, as was demonstrated in a study that identified genes involved in nitrogen-mediated root architecture plasticity in 69 *Arabidopsis* genotypes. This work suggested that the mechanisms controlling roots plasticity in Col-0 are not different from those of other natural genotypes, and Col-0 mutants can be used to mimic the natural variation in a limited population size (Rosas et al., 2013). Recent genomic analysis of pangenomic variation has shown extensive presence/absence variation in genes including enzymes and regulatory genes suggesting the likelihood that other accessions have mechanisms not present in the Col-0 reference (Menz et al., 2018). Thus, we utilized a larger accession collection to survey for novel nitrogen responses/ behaviors that are not possible within Col-0 or its mutants.

To address these questions, we phenotyped growth responses under both nitrate and ammonium to compare if the genetics of natural variation to the two sources are similar or independent. We measured the responses of the genotypes using both developmental phenotyping of the root system architecture and metabolic phenotyping of the defense metabolite GSLs. This allowed us to assess if the two trait classes identify similar or disparate genetic mechanisms and test if the extensive genetic variation in GSLs may contribute to nitrogen responses.

To query for rare accessions with novel responses, we utilized a much larger collection of accessions than previous. This analysis identified a large number of genes involved in the nitrogen response that are predominantly specific to a trait class under a specific nitrogen source under different genetic backgrounds. Having a large collection of genotypes allowed us to identify novel behaviors in a polygenic trait like *Arabidopsis* nitrogen responses. These are likely caused by combinations of diverse alleles and as such would be difficult or impossible to observe in smaller populations or a single reference genotype. Using this vast collection of natural genotypes and studying both developmental and metabolic responses expand our understanding of the processes and mechanisms involved in nitrogen responses.

## Results

### Phenotyping platform optimization

To optimize our ability to measure phenotypic diversity across *Arabidopsis* genotypes to variation in nitrogen, both source and concentration, we quantified seedlings growth growing on a range of nitrate and ammonium concentrations. To maximize the potential trait range measurable across the *Arabidopsis* genotypes we found matching concentrations of the two nitrogen sources that elicited substantial trait variation without constraining growth, e.g., neither an oversupply nor an undersupply. We grew the reference *Arabidopsis* Col-0 on media with either nitrate (NO_3_^−^ as KNO_3_) or ammonium (NH_4_^+^ as NH_4_HCO_3_) as a sole nitrogen source with concentrations ranging from 0.05 mM to 5 mM. After 12 days we measured five developmental traits linked to differential nitrogen responses, including leaf area and root phenotypes (a full list is provided in Figure S1, Tables S1 and S2). As previously reported (Bloom et al. 2012), *Arabidopsis* Col-0 seedlings at ambient CO_2_ grows better with nitrate as a nitrogen source than with ammonium under most concentrations, with higher average weight, larger leaf area, and longer primary roots (Figure S1). On ammonium, *Arabidopsis* Col-0 seedlings performed well from 0.1 mM to 1mM, but the extreme ammonium concentrations (0.05mM and 5mM) prevented seed germination or growth (Figure S1). Therefore, 0.1 mM and 1 mM, the widest pair of concentrations that allowed growth under both nitrogen sources, were chosen as the working concentrations for the experiments to assess genetic variation in *Arabidopsis*.

### Diversity of Arabidopsis genotype responses to ammonium and nitrate measured using developmental and biochemical traits

To measure natural genetic variation in *Arabidopsis* responses to nitrate and ammonium we assessed developmental and defense metabolite traits for a population of 1135 sequenced natural genotypes collected from geographical locations around the world (The 1001 Genomes Consortium, 2016). We included developmental and defense metabolite traits to test if the genetic variation in nitrogen responses had global effects on the *Arabidopsis* genotypes or if each trait may identify different responses and genes.

Each genotype was sown in triplicate on 4 nitrogen conditions: nitrate as a sole nitrogen source in a concentration of 0.1mM or 1mM, or ammonium as a sole nitrogen source in a concentration of 0.1mM or 1mM (for more details see methods). Twelve days after planting, each seedling was measured for developmental and defense metabolite traits (for more details see methods, Tables S3 and S4).

We used a linear model to parse the influence of genotype (genetic variation amongst the genotypes that is independent of nitrogen), environment (trait variation solely influenced by the nitrogen source, and nitrogen source by concentration), and the interaction of genotype and environment (trait variation where genotype variation alters the response to nitrogen variation) on the variation in the developmental traits (Figure 1, sup. Tables 6,7). The linear model also included terms for random effects of experiment and culture plate (for more details see methods). Across all genotypes and nitrogen conditions, genotype (Accession) influenced the majority of explained developmental trait variation. Nitrogen source and the interaction of nitrogen source and concentration significantly altered developmental traits, but these effects were secondary to the interaction of the nitrogen terms (source or concentration) with genotype (Figure 1). This indicates that for developmental traits such as lateral root formation, primary root length, and leaf area, the difference between ammonium and nitrate at these concentrations is less influential than the interaction between nitrogen source or concentration and the genotype. Using the same linear model, we estimated the level of variation due to each of the indicated parameters for each defense metabolite trait. For all the individual defense metabolite traits the variation was determined equally by genotype, and the interaction of nitrogen and genotype (Figure 1). For some of the GSL summation traits (e.g., the C3 vs. C4 ratio = the number of carbons in the GSL backbone, alkenyl ratio, and GSOH activity), genotype explained most of the variation while the interaction of nitrogen and genotype explained a small portion of the variation. This indicates diverse nitrogen responses across genotypes for both the defense metabolite and developmental traits.

**Figure 1:**
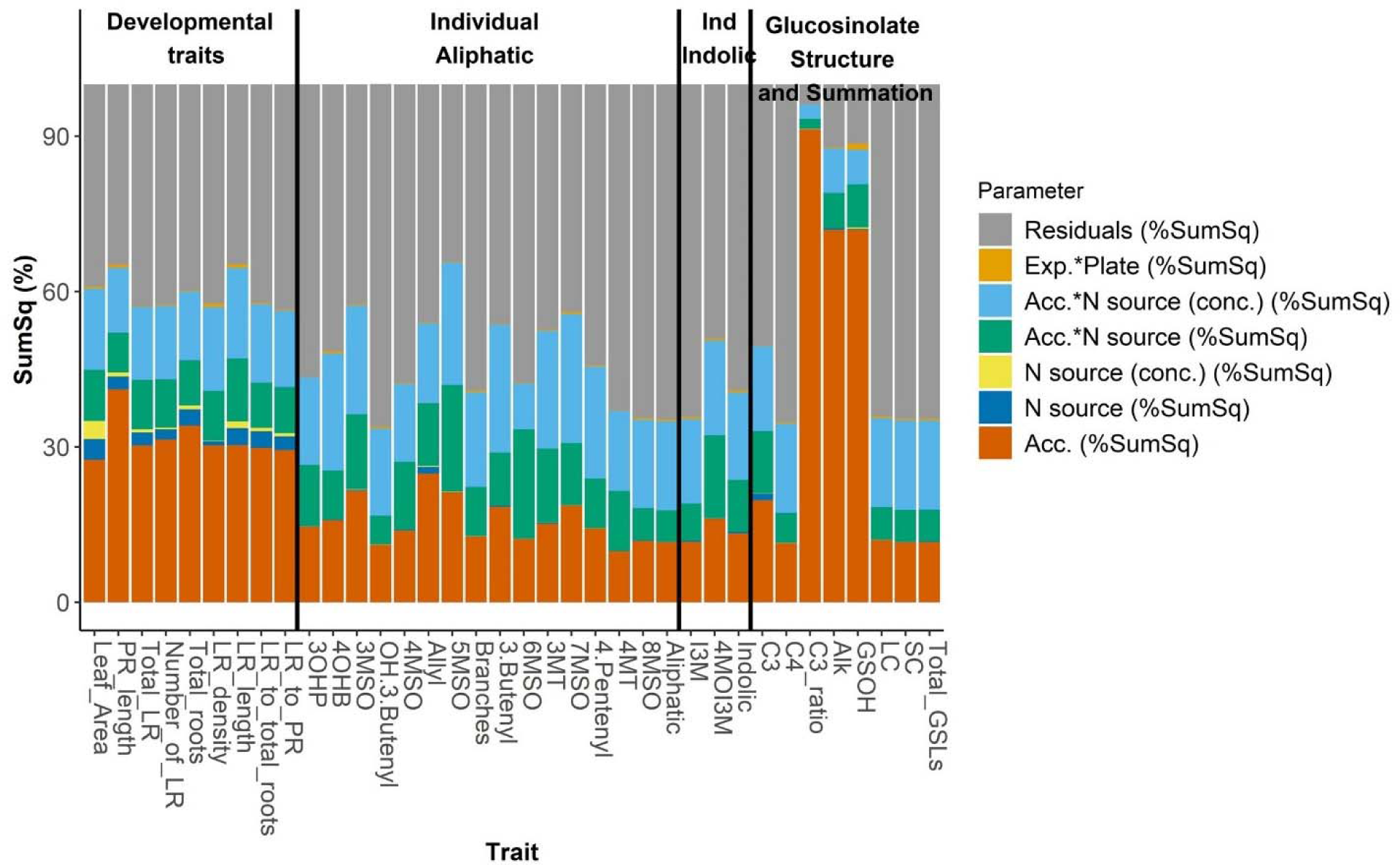
Broad-sense heritability: percentage of variance explained by each variable was estimated for each trait based on a linear model followed by ANOVA with the indicated effects upon the measured trait (Acc= accession, conc=nitrogen concentration, Exp*Plate=experiment block) Ind = individual.

### Genotype and nitrogen interactions create diverse changes in shoot and root development

The linear models indicated a high level of diversity in the genotype’s responses to the four nitrogen conditions. To visualize the potential range of responses across the genotypes, we plotted for the distribution of each trait using each genotype’s adjusted mean phenotypes for each nitrogen condition (Figure 2 A-D).

**Figure 2:**
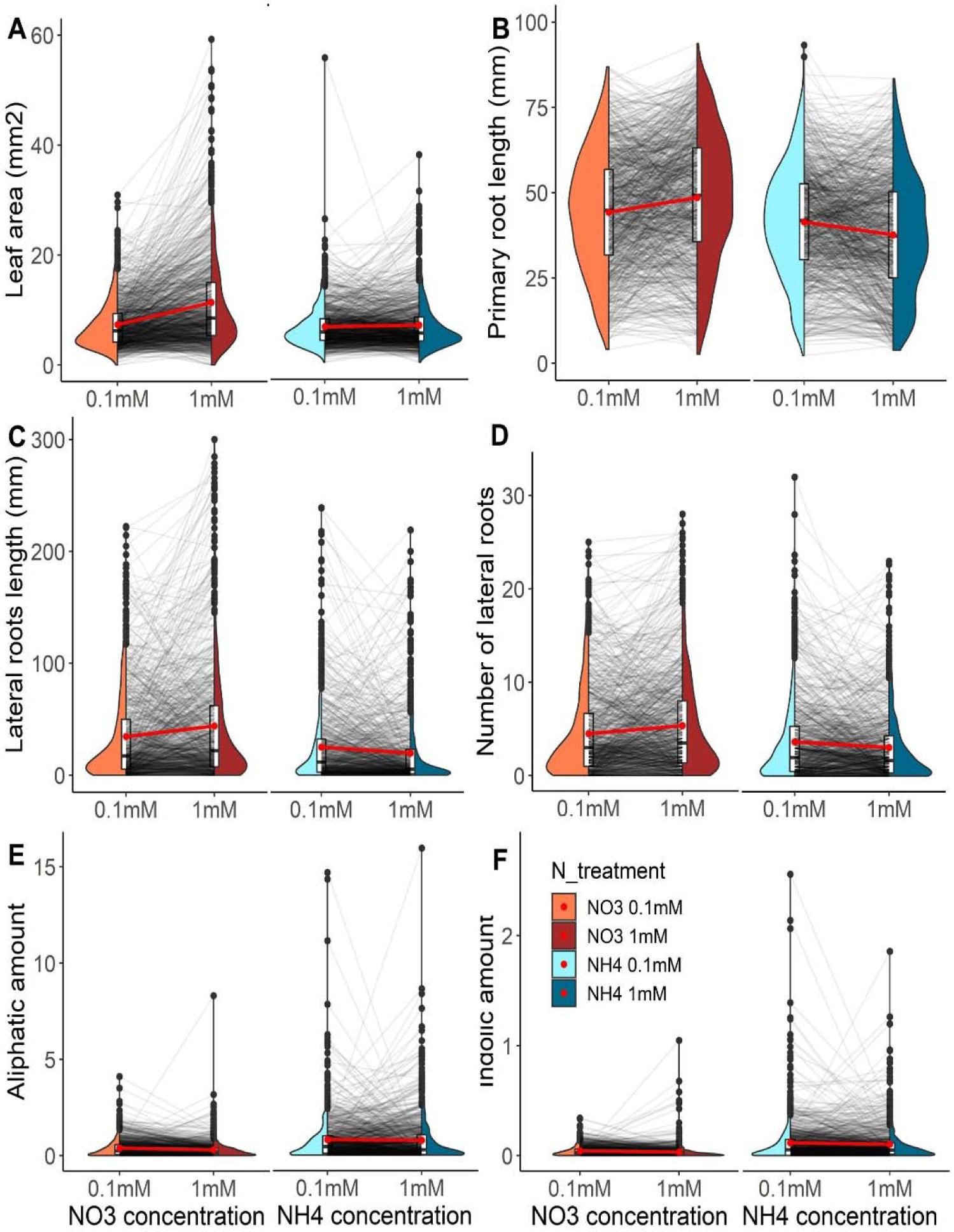
Nitrogen conditions affect different traits across natural accessions: phenotypes of Arabidopsis accessions grown on 4 different nitrogen conditions (nitrate 0.1mM, nitrate 1mM, ammonium 0.1mM, ammonium 1mM). A. Leaf area. B. Primary root length. C. Lateral roots length. D. Number of lateral roots. E. Aliphatic GSLs. F. Indolic GSLs. Grey lines between each pair of nitrogen concentrations connect the means of each individual accession grown under each nitrogen concentration. Red lines between the violins connect the mean phenotype between two concentrations of the same nitrogen source. Aliphatic and indolic GSLs amounts were normalized by total roots length for seedlings grown on NO3 1mM and NH4 1mM (amount units are umol/mm), and by total roots length and leaves area for seedlings grown on NO3 0.1mM and NH4 0.1mM (amount units are umol/mm+mm2).

On average, the population of *Arabidopsis* genotypes had significantly more leaf area, longer primary roots, and more lateral roots on nitrate than ammonium (Figure 2 A-D) (post-hoc TukeyHSD test shown in Table S8). Further, the population average for these traits increased with additional nitrate and decreased with additional ammonium (Figure 2 A-D). Thus, the average *Arabidopsis* genotype tends to present extended growth on nitrate than ammonium and is constrained by nitrate availability, aligning with previous literature (Li et al., 2014; Britto and Kronzucker, 2002; Jian et al., 2018; Fuertes-Mendizábal et al., 2013; Marino and Moran, 2019).

As anticipated from the high fraction of variance attributed to the genotype × nitrogen interactions, the range in the different genotypes’ responses to the nitrogen sources dwarfs the average population response. This includes genotypes that have very different behaviors than the average. For example, while most genotypes present enhanced growth on 1 mM than 0.1 mM nitrate, there are genotypes, including IP-Con-0, IP-Ses-0 and Ven-1, that perform better on the lower nitrate concentration for most of the developmental traits (Figure 2 and Figure S3). These genotypes also tend to perform better on higher ammonium than on higher nitrate. Thus, there is an extensive range of genetically dependent nitrogen responses present amongst these genotypes, including the potential to perform better with ammonium as a nitrogen source. This allows the identification of genotypes that have novel responses in comparison to the Col-0 reference genotype.

### Genotype and nitrogen interactions create diverse defense metabolite responses

To evaluate if defense metabolites exhibit a similar diversity in nitrogen responses, we conducted a similar analysis as above. For this we focused on two defense metabolite traits, the sum of all indolic GSLs (GSLs derived from Tryptophane) and the sum of all aliphatic GSLs (GSLs derived from Methionine) in each seedling, because these traits are quantifiable in all the genotypes (Figure 2 E, F). In contrast, most individual aliphatic GSLs have significant presence/absence variation limiting the ability for direct comparison across the whole population. Plotting the genotype’s adjusted mean phenotypes showed that GSLs are 10-fold higher under ammonium than under nitrate with no significant change across the different nitrogen concentrations (Table S8 and Fig. 2 E,F). This agrees with previous publications, and was suggested to be a strategy to avoid ammonium toxicity by diverting ammonium to GSL production (Marino and Moran, 2019; La, 2013; Marino et al., 2016; Coleto et al., 2017). This pattern differs from the developmental traits suggesting that GSLs have different nitrogen responses.

Defense metabolite traits, like the developmental traits, display a large diversity of genotype by nitrogen interactions (Figure 2 E,F). This is supported by the linear model showing that the genotype by nitrogen interaction explained most of the trait variation. While the average genotype showed minimal response to the different nitrogen concentrations under both nitrogen sources, certain genotypes exhibited enhanced nitrogen sensitivity. Both the IP-Ara-4 and IP-Mah-6 genotypes accumulated significantly higher amounts of both aliphatic and indolic GSLs when grown on the higher concentration of ammonium (Figure S3, Table S9). In contrast, IP-Ara-4 seemed to decrease GSLs (aliphatics and indolics) when grown on higher nitrate. This further demonstrates the potential of this collection of genotypes to identify novel nitrogen responses that differ from the average genotype or from the Col-0 reference genotype.

### Partial correlation between developmental and metabolic traits

Because mechanistic links connecting nitrogen, development, and defense metabolism are possible, we tested for such links in the population. We calculated genetic correlations between the traits under the assumption that mechanistic links would show up as shared genetic causality and result in correlated trait variation. To minimize the influence of any nitrogen main effect on the analysis we correlated across each nitrogen condition, source or concentration, separately. This showed that all the developmental traits are significantly positively correlated (Figure 3). In contrast, the defense metabolites showed more diversity in their correlations both within and across nitrogen conditions (Figure 3). For example, the amount of indolic GSLs is significantly positively correlated with all the developmental traits when the genotypes were grown on 0.1 mM nitrate or 1 mM ammonium. However, this correlation is largely non-existent when the accessions were grown on 1 mM nitrate or 0.1 mM ammonium. In contrast, the aliphatic GSLs amount shows a low correlation to developmental traits when the genotypes were grown on 1 mM nitrate or 0.1 mM ammonium, but higher correlation when the genotypes were grown on 0.1 mM nitrate or 1 mM ammonium. This suggests that there is some shared genetic causality across the developmental and defense metabolite traits but that it is highly conditional to the specific nitrogen condition.

**Figure 3:**
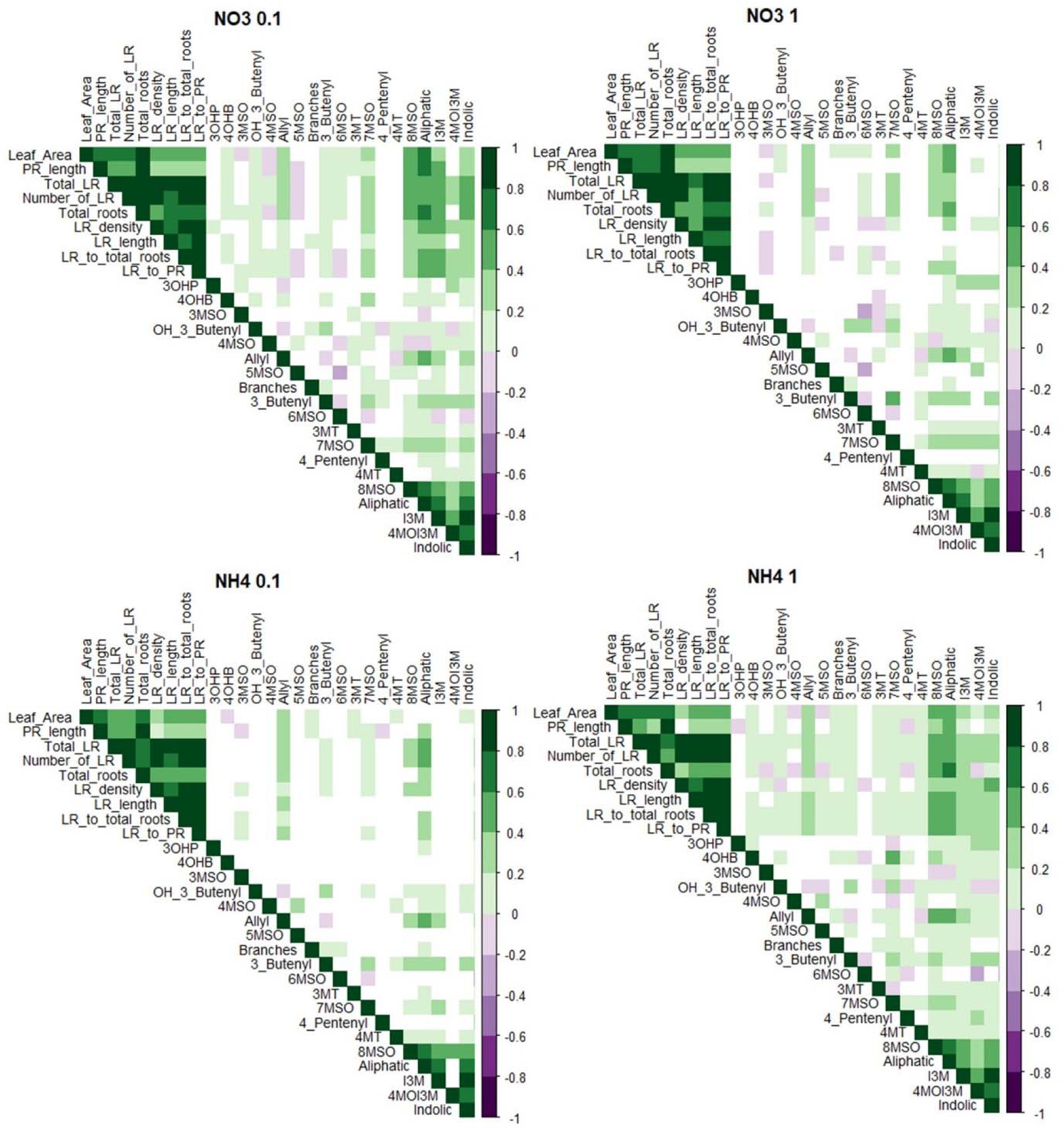
Nitrogen conditions present different correlations between the traits: correlation matrix between the traits under the different nitrogen conditions. Colors represent the correlation values. White squares represent nonsignificant correlations (method: Spearman, conf.level = .95).

The combination of developmental and defense metabolite traits also allows us to determine if there is modularity amongst the genotypes whereby groups of genotypes show similar responses to the nitrogen conditions. If the *Arabidopsis* genotypes are evolving to a limited range of nitrogen conditions that are distinct from each other, a modular structure might be expected to reflect this structure. The alternative option is that nitrogen responses exist on continuums with no grouping in the genotypes. To test these possibilities, we took the developmental and defense metabolite traits across all four nitrogen conditions and used principal component analysis (PCA) to assess the structure of the phenotypic variation. The PCA showed that there was no readily identifiable genotype grouping and that phenotypic variation in genotypes exist on a multi-dimensional continuum. Further, the PCA largely did not separate by nitrogen conditions supporting the importance of genotype × nitrogen interactions (Figure S7).

### Candidate genes associated with nitrogen responses

To identify loci underlying genetic variation in these nitrogen responses, we performed genome-wide association studies (GWAS, with EMMAX algorithms) using the dense SNP data available for this genotype collection. We focused on four developmental traits (leaf area, primary root length, lateral roots length, and number of lateral roots) and the two defense metabolite traits (aliphatic and indolic) under each nitrogen condition as traits for GWAS. We also included the log (10) ratio for each trait in each genotype across the two concentrations of the same nitrogen source as additional traits to approximate the nitrogen responsivity (Figure S8). This resulted in 36 different GWAS: six traits across four nitrogen conditions and two ratios per the six traits (Figure S9). Each trait yielded a diverse group of significant SNPs (we observed low overlap between SNPs from each GWAS, Figures S10 – S13). Amongst this dataset, no large effect loci were detected which suggested a poly- or oligo-genic architecture.

To filter for core genes and other trends, we combined each group of traits (development, aliphatic, and indolic) under each nitrogen source as a set (Figures S10 – 13, all future references of “set” refer to those groups of traits under each nitrogen source). We combined leaf and roots traits into one set of developmental traits because they show a high correlation under all nitrogen conditions (Figure 3), and previous mechanistic work showing that these traits can have a coordinate genetic control; for example SnRK (SNF1-related protein kinase) gene family control both roots and shoots growth through the activation of abscisic acid (Fujii and Zhu, 2009), and TOR (TARGET OF RAPAMYCIN) kinase (McCready et al., 2020). For each of these sets, we identified the genes that have the most consistent effect across all individual GWA within a set using the Multivariate Adaptive Shrinkage (MASH) method. Then the MASH method uses all the SNPs in the set and ranks them based on their shared and consistent effects across the set. This rank is then identified the SNPs with the most consistent effect on traits to shrink the number of SNPs/genes for further interrogation (for details see methods). For example, we took all of the EMMAX output for GWA for all the developmental traits on all the nitrate conditions (12 total trait GWA datasets), combined them to a set, and used MASH to find the shared SNPs that influence development across nitrate conditions. This analysis still found 1281 genes as being at the core of this set of GWAS.

Testing for overlap between the six gene lists from the six trait sets from the MASH highlighted candidate genes that were mutual to the nitrogen sources and/or trait types (Figure S16). Overall, around 30% of the genes in each set were found in at least one other set (Table S11). This indicates that most of the genes in most of the sets are specific to one set and are not mutual across other nitrogen sources and/or traits (Figure 4, Table S11). We then tested for an overlap between genes identified using the GWA on the same trait type across the different nitrogen sources. This test for more shared causality within the developmental traits across nitrate and ammonium. Conducting Fisher’s exact test to measure overlap of genes commonly found from different sets shows significant enrichment for common genes influencing developmental traits across nitrate and ammonium (Figure 4). In contrast, both defense metabolite traits had no evidence for enrichment across the two nitrogen sources, suggesting that largely different genetic networks may influence defense metabolism across the nitrogen sources. Next, we shifted the analysis to test for overlap between trait sets on a specific nitrogen source; for example, did ammonium have more commonality across aliphatic and indolic GLS. This showed that there was a significant enrichment in gene overlap in the indolic and aliphatic GLS trait sets under ammonium conditions, but almost none in the sets under nitrate conditions. This suggests that when seedlings are grown on ammonium, shared genes influence variation in both the indolic and aliphatic GLS pathways (Figure 4, and Table S12).

**Figure 4:**
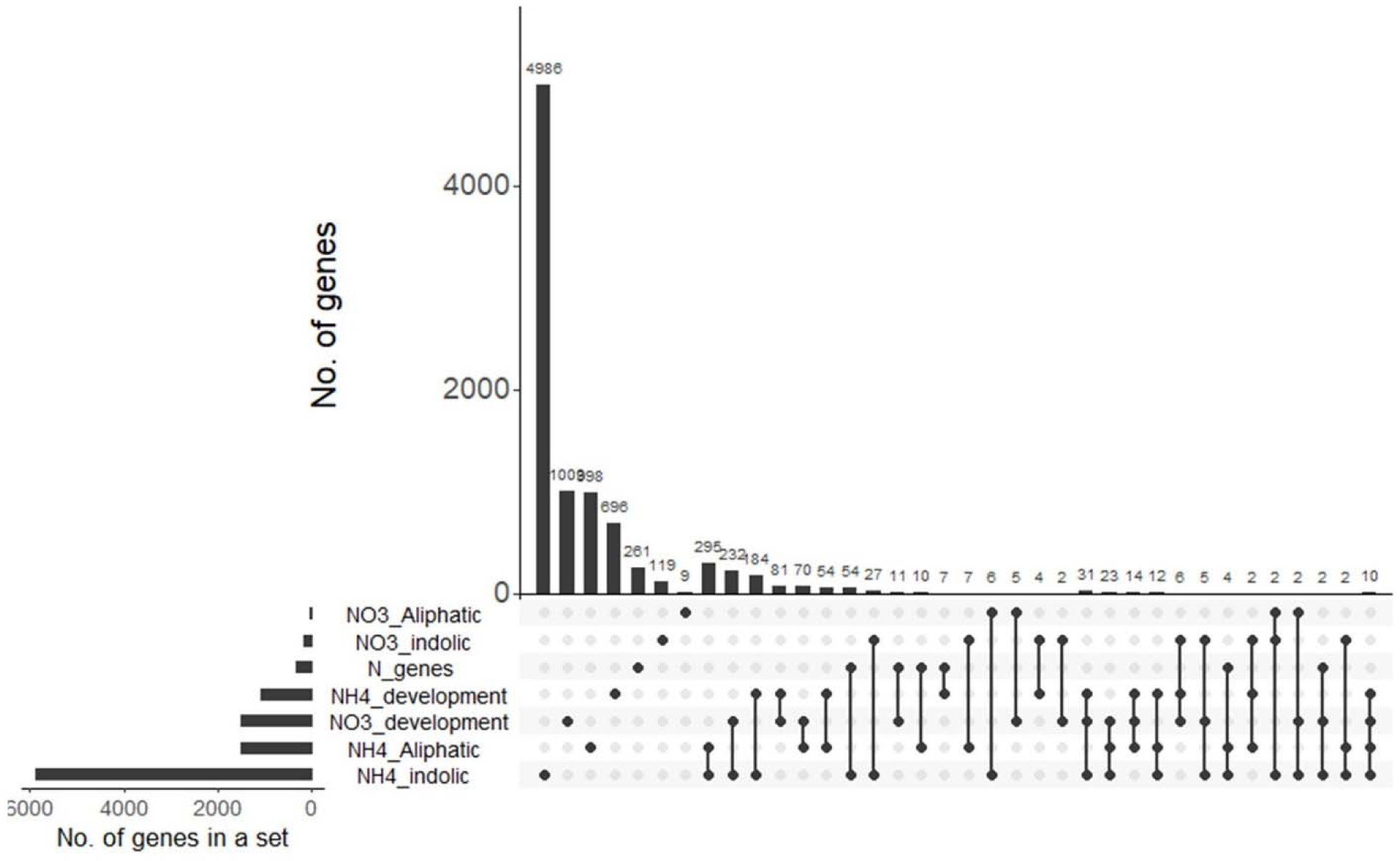
Overlaps between genes yield from the different mash sets: SNPs with mash score > 0.98 were transformed to genes, and the overlap of these genes between the different sets was analyzed. N genes are a set of Arabidopsis genes known to be related to some nitrogen process. Fisher’s exact test was performed for each pair of sets (supp. Table 12). Total number of genes = 28496.

We then assessed whether the candidate genes we identified were already known to be related to some nitrogen process (assimilation, signaling, transport etc.) or whether inclusion of defense metabolites and ammonium identified new possible nitrogen-associated genes. *Arabidopsis* has a large catalog of genes with previously ascertained mechanistic links to nitrogen, typically derived using root traits and nitrate. Using literature annotation, we created a set of 302 genes that are known in Arabidopsis to be related to some nitrogen process (Alvarez et al., 2014; Brooks et al., 2019; Castaings et al., 2009; Cheng et al., 2021; Gaudinier et al., 2018; Gifford et al., 2008; Konishi and Yanagisawa, 2013; Krouk et al., 2010; Medici et al., 2015; Obertello et al., 2010; Rubin et al., 2009; Varala et al., 2018; Vidal et al., 2014; Xu et al., 2 016; Zhang and Forde, 1998) (For details see methods, and Table S13). Using this list of known mechanistic genes, we checked for an overlap with the six gene sets from the MASH analysis (Figure 4). Among the genes that appeared in more than one set and that are known to be involved in some nitrogen response are genes involved in development (AT4G18390, TEOSINTE BRANCHED 1), nitrate regulation (AT3G60320) and a nitrate reductase (AT2G15620), and hormone regulation (AT5G51190 is a member of the ERF/AP2 domain transcription family, and AT4G39070; BZS1 that is regulated by brassinosteroids through BZR1). Gene Ontology (GO) analysis with the full list of genes that showed up in 3 sets or more did not yield any enrichment.

A test of the scope of overlap between the 6 sets and the known nitrogen genes showed that less than 1% of the genes in each trait set overlapped with the set of known nitrogen genes, regardless of if nitrate or ammonium was the nitrogen source. Fisher’s exact test indicates that this overlap is non-significant indicating that known nitrogen associated genes are not enriched (Table S12). Therefore, most of the candidate genes were not previously linked to nitrogen responses in *Arabidopsis*.

To obtain insights about the biological processes in which the associated candidate genes are involved, we conducted GO analysis (Figure S16). Analyzing each MASH derived gene list for the six trait sets, showed some high-level GO terms that were enriched, like metabolic processes and defense responses. However, these were only higher-level GO terms with no especially informative connections (Table S15). To test for a more limited gene set common across a few trait sets we used genes that appeared in more than one sets. Using both the developmental and metabolic trait sets, the biological processes that are enriched based on these genes, include both GSL and general nitrogen metabolism, and categories around cell processes (Table S14).

### Candidate nitrogen-related genes are co-expressed

Previous work has shown that co-expression modules of candidate GWAS genes can identify genes that are more likely to be casually connected to the traits of interest (Chan et al., 2011; Wisecaver et al., 2017). To test if co-expression filtering of the candidate genes may identify more mechanistically insightful gene sets as previously suggested, we developed co-expression gene modules using expression data from seedlings of all the genotypes (Figure S16) (Kawakatsu et al., 2016; Wisecaver et al., 2017). Previous work has shown that co-expression modules do not have to be derived from transcriptomic experiments designed around the specific conditions being tested and that any transcriptomic experiment can provide highly informative module. This extends even to the observation that that informative co-expression modules can be readily obtained from stochastic inter-individual variation within a single genotype (Liu et al., 2021). This analysis resulted in 2864 modules with a median of 7 genes in a co-expression module (range 3 to 257 genes). Of these co-expression modules, 436 had no GWA candidates.

Using these co-expression modules, we tested if our GWA candidates clustered into specific co-expression modules. We assigned each gene in each set to a co-expression module(s), thereby creating a list of modules for each set, based on the genes in each set. Candidate genes not assigned to a co-expression module were discarded. By filtering for likely causal genes with these co-expression modules, we found more overlap across gene modules of trait sets than previously found using the original gene lists (Figure S17). Whereas the simple gene lists found little overlap between traits sets, the co-expression modules are highly shared between the trait sets: 90% of the identified co-expression modules were associated with at least 2 different trait sets, and only 10% of the modules were specific to one trait set (Table S11). For example, one co-expression module (module 104) was found to be associated with candidate genes from all trait sets, suggesting that this module of 119 genes may play a common role in nitrogen responses across development and defense. To learn about the biological processes associated with the modules that were associated with at least five of the trait sets (24 modules, Table S16), we used the genes in those modules in a GO enrichment analysis. This showed that these co-expression modules associated with most nitrogen traits are enriched for genes involved in root development and cell division (Table S17). These processes are expected to be involved in altering plant development and metabolism. This means that while most of the candidate genes show specificity to distinct nitrogen responses (a specific set), they appear to relate to a few commonly identified modules, as previously suggested (Gaudinier et al., 2018). This suggests that the natural variation may be functioning through a defined set of modules that can be found using the combined GWA/co-expression approach.

## Discussion

How a plant uses nitrogen is critical to the plant’s productivity and fitness, especially in a changing environment. Developing a better understanding of the genetics and mechanisms that govern plant nitrogen use will facilitate engineering or breeding plants to use nitrogen more efficiently. To map the potential diversity of processes involved in plant nitrogen response and use, we grew a large population of *Arabidopsis* genotypes under four different nitrogen environments, and measured a set of developmental traits, and a set of defense metabolite traits. We found that this population presents a diversity of responses to the different nitrogen conditions associated with a large number of genes. While most of those genes are specific to distinct nitrogen conditions (a particular nitrogen source or concentration), it is possible to find genes involved in potentially responding across nitrogen conditions using a combined GWA/co-expression filtering method.

### Large population size allows the identification of rare and novel nitrogen responses amongst the genotypes

Using a large number of genotypes, we found that the population had a clear average response while individual genotypes had a wide range of responses to nitrogen across the different traits. The large population size allowed us to identify extended phenotypic tails that include previously unobserved responses of *Arabidopsis* to nitrogen. For example, while most of the genotypes presented elevated root and shoot growth on nitrate over ammonium, several genotypes presented the opposite behavior with increased growth when grown on ammonium. Extended phenotypic tails in natural populations can be caused by variation in either a few major loci or multiple small effect loci. Several lines of evidence suggest that we observed mainly variation in small effect genes that blend to create these unique events. First, most of the traits show a unimodal distribution of phenotypes with a long continuous tail of phenotypic variation and lack any evidence of multi-modality or extreme outliers (Schraiber and Landis, 2015). These distributions imply that these traits are controlled by different combinations of small effects alleles.

The second line of evidence that mainly small effect genes control these traits in the population comes from the GWAS we conducted using the different traits under the different nitrogen environments. The GWAS did not find evidence of any large-effect loci associated with the responses to nitrogen. Instead, the loci were small effects, suggesting that the unusual phenotypic responses arise from combinations of alleles at a large number of loci. The potential for novel phenotypes to arise by accumulation of alleles at multiple loci within a network was recently presented by a model showing that trait values can be non-linearly created by a network of interactions with different weights (Milocco and Salazar-Ciudad, 2022). To check for potential rare large-effect loci we queried the genotypes at the distribution tails for a preponderance of large-effect natural knockouts in known nitrogen genes but were unable to find any overlap.

Together these results suggest that using a large population of genotypes with natural variation is a preferable strategy than using a single reference genotype, e.g., Col-0, because it improves the detection of responses that are a result of different combinations of multiple alleles. Therefore, using a vast collection of natural genotypes and combining both genomic and phenotypic techniques can be a good strategy to study complex traits, such as the response to nitrogen conditions. Further, these unique phenotypes (e.g., those that present extensive growth on ammonium than nitrate) may be generated by stacking small effect loci to create a large phenotypic variance.

### Metabolic and Defense metabolites provide different readouts of the plants’ nitrogen response

To test if the genetic basis of nitrogen responses is uniform across a range of traits or may be somewhat trait specific, we analyzed two sets of traits that are associated with nitrogen responses. First, we used classical developmental traits that include shoot and root descriptors associated with nitrogen responses. We also measured the response of defense metabolites that include two families of GSLs, indolic and aliphatic, that depend on amino acid availability for their synthesis. Together we used these two sets of responses to compare the effects of nitrogen on different processes in *Arabidopsis* seedlings. The phenotypic analyses revealed that each set of traits presented different patterns of nitrogen responses and preferences across the population. Further, the GWAS based on these two sets of traits yielded loci that are associated with each set of traits, and a set of loci that is shared between the traits. This demonstrates the importance of studying different types of responses (e.g., metabolic responses) under the same conditions when studying complex responses because each set of traits highlights different aspects in the plants response to nitrogen and can potentially identify different genetic mechanisms involved in nitrogen responses.

### Nitrate and ammonium induce different responses in plants

To test if there is a universal nitrogen response in *Arabidopsis* or if nitrate and ammonium may identify different genetic mechanisms, we measured trait variation using both nitrate and ammonium as a sole nitrogen source. Conducting GWAS using traits of genotypes that grew on each of these nitrogen sources revealed two sets of loci, each one of which is associated with one of the nitrogen sources, and a set of loci that are shared among the two nitrogen sources. Most of the genes in this analysis are specific to one nitrogen source (68% of the nitrate related genes, and 93% of the ammonium related genes), and only a small portion of the genes is shared across the two sources. Analyzing the correlations between the different traits revealed that while the developmental traits presented strong correlations to each other across all the nitrogen conditions, the GSLs traits presented different patterns of correlations between each other and the developmental traits, depending on the nitrogen condition provided to the plants. Together, these results suggest that while there is a shared set of genes that will influence the plant across all nitrogen sources, there is an additional, potentially larger, set of genes that will be specific to the nitrogen condition (source and concentration) that the plants experience. Therefore, the exact nitrogen condition (source and concentration) has a dramatic effect on the genes and mechanisms used by the plant to optimize its performance.

### New gene identification

Another benefit of combining multiple traits, multiple environments, and a larger population is the potential to identify a large collection of potential candidate nitrogen response genes. Interestingly, the overwhelming majority of these candidate genes have not been previously associated with nitrogen responses. However, many are associated with expected mechanisms such as root development, cell-cycle regulation, and metabolic processes. The new genes that these analyses identified introduce new players potentially involved in plant response to nitrogen. Future validation efforts of these genes will help to provide a better understanding of the mechanisms shaping a plant’s response to a specific nitrogen source. Overall, our experiment suggests that using such a large population of genotypes and analyzing diverse responses under different environments can reveal new genes that are potentially associated with nitrogen responses.

## Acknowledgements

We thank Dr. Alice MacQueen and Dr. Thomas Juenger (The University of Texas, Austin) for their guidance and discussion on MASH analyses, Dr. Dan Runcie (Department of Plant Sciences, University of California, Davis) for discussion on model development, Dr. Allison Gaudinier (Department of Plant and Microbial Biology, University of California, Berkeley), Elli Pana Cryan (the Department of Plant Sciences and the Department of Evolution and Ecology, University of California, Davis) and Paul Kasemsap (Department of Plant Sciences, University of California Davis, Davis, CA, USA) for critical reading of the manuscript, and Heera Rasheed, Jozanne Berdan, Janine Tsai, and Bryce Gregg for assistance in the experiments. This work was supported by the National Science Foundation, Directorate for Biological Sciences, Division of Molecular and Cellular Biosciences (grant no. MCB 1906486 to DJK) and Division of Integrative Organismal Systems (grant no. IOS 1655810 to DJK and AJB).

## Methods

### Plant material and experimental design

Seeds for 1135 *Arabidopsi*s (*A. thaliana*) genotypes were obtained from the 1001 genomes catalog of *A. thaliana* genetic variation (The 1001 Genomes Consortium, 2016); https://1001genomes.org/).

Seeds were surface sterilized with 4.125% (v/v) sodium hypochlorite and 0.01% (w/v) Tween 20 for 15 min, then rinsed five times with distilled water.

The seeds were next cultivated on 13 CM^2^ plastic culture plates containing the following: 0.1 mM or 1 mM KNO_3_ or NH_4_HCO_3_, 4 mM MgSO_4_, 2 mM KH_2_PO_4_, 1 mM CaCl_2_, 10 mM KCl, 36 mg/l FeEDTA, 0.146 g/l 2-morpholinoethane sulfonic acid, 1.43 mg/l H_2_BO_3_, 0.905 mg/l MnCl_2_·4H_2_O, 0.055 mg/l Zinc sulfate heptahydrate, 0.025 mg/l copper sulfate, 0.0125 Na_2_MoO_4_·2H_2_O, 1% sucrose, 0.8% (w/v) agar, pH 5.7. The plates were placed vertically, and the seedlings were grown at 22°C/24°C (day/night) under long-day conditions (16 hr of light/8 hr of dark). The petri dishes with the seeds were then placed at 4°C at the dart for 3 days. Germination timing was homogenous across the collection, and seedlings were collected 12 days following planting. For the initial concentration selection experiments, additional concentrations of KNO_3_ and NH_4_HCO_3_ were analyzed.

Each genotype was sowed on all four N conditions with three replicates per condition. The replicates were sowed on separate plates. Each plate contained six different seeds. Each experimental block contained 60 different genotypes and was referred as “Experiment”.

### Roots and leaf measurements

The developmental traits that were measured to describe the plants response to the different nitrogen conditions are leaf area, primary root length, total lateral roots length (the sum of the length of all the lateral roots in a seedling) and the number of lateral roots. Root traits were measured using Rootnav (Pound et al., 2013) or ImageJ software (https://imagej.nih.gov/ij/) when the roots were tangled with each other.

Leaf area was measured using an automated image processing workflow developed in Python, using functions from OpenCV and PlantCV packages (Gehan et al., 2017), with the following steps: cropping agar plate region, leaf identification, pixel counting, and scale identification. *GSL extractions and analyses:* To compare metabolic and developmental responses, the seedlings were collected and assayed for the presence and amounts of individual glucosinolates (GSLs) from two families based on the amino acid from which they derived: indolic GSLs derived from tryptophan, and aliphatic GSLs derived from methionine. As the individual GSLs are derived from amino acids, the sums of the GSLs provide a sensitive indication of the availability of these amino acids within the plant, and an indication of how they are affected by the nitrogen condition provided to the seedling. We measured 2 individual indolic GSLs, 15 individuals aliphatic GSLs, and calculated several traits that derived from those individual GSLs (Table S6, S7). GSLs were measured as previously described (Kliebenstein et al., 2001a; Kliebenstein et al., 2001b; Kliebenstein et al., 2001c). Briefly, each seed was harvested in 200 μL of 90% methanol. Samples were homogenized for 3 min in a paint shaker, centrifuged, and the supernatants were transferred to a 96-well filter plate with DEAE sephadex. The filter plate with DEAE sephadex was washed with water, 90% methanol and water again. The sephadex-bound GSLs were eluted after an overnight incubation with 110 μL of sulfatase. Individual desulfo-GSLs within each sample were separated and detected by HPLC-DAD, identified, quantified by comparison to standard curves from purified compounds A list of GSLs and their structure is given in table S5. Raw GSLs data are given in table S3. GSLs ratios were calculated as following: C3 ratio=C3/(C3+C4), Alk=(OH-3-Butenyl+3-Butenyl+Allyl)/(3OHP+3MSO+Allyl+3MT+OH-3-Butenyl+4MSO+3-Butenyl+4MT+4OHB), GSOH=OH-3-Butenyl/(OH-3-Butenyl+3-Butenyl).

### GSLs normalization

GSL amounts were normalized for each nitrogen condition separately, based on leaf area and total roots length. For each nitrogen condition, a linear model was utilized to analyze the effect of leaf area and the total roots length upon the seedlings weight. The Intercept and the slope of leaf area and the total roots length were used to normalize the GSL amounts. This was to provide a common unit of comparison when it was not possible to measure the mass of all the seedlings.

For seedlings grown on KNO_3_ 0.1mM: 9.902e-06×(total roots length) + 7.706e-05(leaves area) + 1.169e-03.

For seedlings grown on KNO_3_ 1mM: 8.601e-05×(total roots length) - 1.313e-03.

For seedlings grown on NH_4_HCO_3_ 0.1mM: 6.352e-06 × (total roots length) + 1.491e-04(leaves area) - 1.528e-04.

For seedlings grown on NH_4_HCO_3_ 1mM: 4.768e-06×(roots length) + 1.124e-03.

### Statistics, heritability, and data visualization

Statistical analyses were conducted using R software (https://www.R-project.org/) with the RStudio interface (http://www.rstudio.com/). For each trait, a linear model followed by ANOVA was utilized to analyze the effect of genotype, nitrogen condition, the experiment, and the culture plate upon the measured trait (Trait ∼ Accession+ N source+ (N source/N concentration) + Accession *N source+ Accession *(N source/N concentration) + (Experiment/Culture plate)). In these models the nitrogen concentration is nested within the nitrogen source. Models where the nitrogen concentration is not nested within the nitrogen source were analyzed as well and provided similar outcomes. Broad-sense heritability (Table S6) for the different metabolites was estimated from this model by taking the variance due to genotype (Accession) and dividing it by the total variance. Estimated marginal means (emmeans) for each accession were calculated for each trait based on the genotype and nitrogen treatment using the package emmeans (CRAN - Package emmeans) (Table S4). Differences between pairs of treatments were estimated for each trait using a TukeyHSD test. Data analyses and visualization were done using R software with tidyverse (Wickham et al., 2019) and ggplot2 (Kahle and Wickham, 2013) packages. Fisher exact test was calculated using the library “stats”.

PCAs were done with FactoMineR and factoextra packages (Abdi and Williams, 2010).

Correlations were done using the corrplot package (Wei T, Simko V (2021). R package ‘corrplot’: Visualization of a Correlation Matrix. (Version 0.90), https://github.com/taiyun/corrplot.), based on Spearman correlation.

### Genome-wide association studies

For each trait, GWA was implemented with the easyGWAS tool (Grimm et al., 2017) using the EMMAX algorithms (Kang et al., 2010) and a minor allele frequency (MAF) cutoff of 5%. The results were visualized as Manhattan plots using the qqman package in R (Turner, 2014).

### Multivariate adaptive shrinkage

To reduce the number of SNPs in each set we chose the strongest 100,000 SNPs in each GWA, which included all the significant SNPs in a GWA, and other SNPs. To combine the GWA analysis, MASH was conducted using the mashr package in R (Urbut et al., 2019). SNP’s P-values and standard errors produced by GWAS were used in the MASH analyzes as Bhat and Shat. A subset of 50K SNPs was used as a ‘random’ set to learn the correlation structure among null tests and to fit the mashr model. The analyzes was done with a subset of the strongest 100K SNPs across each set, that was chosen in two ways: 1. Choosing 50K SNPs with the maximum -log10(p-values), by that accounting for strong effects that are specific to traits/ concentration (specific to each GWA). 2. Summing the -log10(p-values) across the set for each SNP and chose the 50K SNPs with the maximum values. SNPs with local false sign rates (lfsr) values higher than 0.98 were chosen to further analyzes.

The SNPs were connected to genes using a window of 2,000 bp upstream of the transcription start site to 2,000 bp downstream of the stop codon.

### Filtering SNPs from the six sets

SNPs from GWA analyses were grouped into six sets, based on the traits and the nitrogen source upon which it was grown: 1. Developmental traits grown on nitrate. 2. Developmental traits grown on ammonium. 3. Aliphatic GSLs grown on nitrate. 4. Aliphatic GSLs grown on ammonium. 5. Indolic GSLs grown on nitrate. 6. Indolic GSLs grown on ammonium (Figures S10-13). Each set includes the traits grown under the two nitrate concentrations, and the logged ratio of the concentrations of each trait under the same nitrogen source.

To analyze the SNPs in each of the six sets we used multivariate adaptive shrinkage (MASH) method and estimated the effect of SNPs in each set of GWAS. Using this method, we could estimate the effect sizes of both shared and condition-specific effects within each one of our six sets (Urbut et al., 2019). This enabled us to detect genes (based on the detected SNPs) that are associated with specific traits (e.g., developmental/ aliphatic/ indolic), genes that are associated with specific nitrogen source (nitrate or ammonium), and genes that are shared across the different nitrogen conditions and/or traits. For each set we conducted the MASH analyses by considering both specific effects (trait specific and concentration specific) and mutual effects across all the GWAS that are in the same set (for more details, see below). Using this method, we were able to rank the SNPs based of their effect in each set and provide for each SNP a score ranging from 0 to 1, where 1 represents the strongest effect. We used these MASH scores of each SNP from each set to choose SNPs with the highest effect from each set. Here we chose SNPs with MASH score higher than 0.98 (Figure S14). Plotting those SNPs based on their location on the chromosome showed that the chosen SNPs from each set are spread across the genome, and we could not detect specific area/s with high SNPs density (Figure S15). We then transformed those SNPs to genes considering 2K bps before and after the start and end point of each gene. This analyzes yield six sets of genes, each set is associated with a different set of traits under a specific nitrogen source (Table S10) and contain between 22 to 5051 genes.

### List of nitrogen related genes in Arabidopsis

Using several literature sources we created a set of genes that have been reported to be associated with some nitrogen process (Table S13). These processes include nitrogen assimilation, transport, metabolism, signaling, and transcription factors (Alvarez et al., 2014; Brooks et al., 2019; Castaings et al., 2009; Cheng et al., 2021; Gaudinier et al., 2018; Gifford et al., 2008; Konishi and Yanagisawa, 2013; Krouk et al., 2010; Medici et al., 2015; Obertello et al., 2010; Rubin et al., 2009; Varala et al., 2018; Vidal et al., 2014; Xu et al., 2 016; Zhang and Forde, 1998).

### Gene ontology enrichment

Gene ontology enrichments were tested using TAIR (https://www.arabidopsis.org/tools/go_term_enrichment.jsp), and with agriGO (Du et al., 2010).

### Co-expression modules

Co-expressed gene sets were identified using the mr2mods program and previous data measuring 874 transcriptomes from natural Arabidopsis genotypes was utilized (Kawakatsu et al., 2016; Wisecaver et al., 2017). Briefly, these packages take the data and creates and all by all Spearman correlation coefficient matrix for the complete transcriptomes. This matrix is then converted into mutual ranks for each transcript pairs. Module groupings are then defined using the mutual ranks as converted to edge weights using an exponential decay function. 8 different exponential decay values ranging from 2 to 100 were evaluated and a decay value of 50 was chosen to maximize average module size.

## Parsed Citations

Abdi, H. and Williams, L. J. (2010). Principal component analysis. WIREs Comp Stat 2, 433–459.

Alvarez, J. M., Riveras, E., Vidal, E. A., Gras, D. E., Contreras-López, O., Tamayo, K. P., Aceituno, F., Gómez, I., Ruffel, S., Lejay, L., et al. (2014). Systems approach identifies TGA1 and TGA4 transcription factors as important regulatory components of the nitrate response of Arabidopsis thaliana roots. Plant J. 80, 1–13.

Bakker, E. G., Traw, M. B., Toomajian, C., Kreitman, M. and Bergelson, J. (2008). Low levels of polymorphism in genes that control the activation of defense response in Arabidopsis thaliana. Genetics 178, 2031–2043.

Beekwilder, J., van Leeuwen, W., van Dam, N. M., Bertossi, M., Grandi, V., Mizzi, L., Soloviev, M., Szabados, L., Molthoff, J. W., Schipper, B., et al. (2008). The impact of the absence of aliphatic glucosinolates on insect herbivory in Arabidopsis. PLoS ONE 3, e2068.

Benderoth, M., Textor, S., Windsor, A. J., Mitchell-Olds, T., Gershenzon, J. and Kroymann, J. (2006). Positive selection driving diversification in plant secondary metabolism. Proc Natl Acad Sci USA 103, 9118–9123.

Bloom, A. J. (2015). The increasing importance of distinguishing among plant nitrogen sources. Curr. Opin. Plant Biol. 25, 10–16.

Bloom, A. J., Burger, M., Rubio Asensio, J. S. and Cousins, A. B. (2010). Carbon dioxide enrichment inhibits nitrate assimilation in wheat and Arabidopsis. Science 328, 899–903.

Bloom, A. J., Asensio, J. S. R., Randall, L., Rachmilevitch, S., Cousins, A. B. and Carlisle, E. A. (2012). CO2 enrichment inhibits shoot nitrate assimilation in C3 but not C4 plants and slows growth under nitrate in C3 plants. Ecology 93, 355–367.

Brachi, B., Meyer, C. G., Villoutreix, R., Platt, A., Morton, T. C., Roux, F. and Bergelson, J. (2015). Coselected genes determine adaptive variation in herbivore resistance throughout the native range of Arabidopsis thaliana. Proc Natl Acad Sci USA 112, 4032–4037.

Britto, D. T. and Kronzucker, H. J. (2002). NH4+ toxicity in higher plants: a critical review. J. Plant Physiol. 159, 567–584.

Brooks, M. D., Cirrone, J., Pasquino, A. V., Alvarez, J. M., Swift, J., Mittal, S., Juang, C.-L., Varala, K., Gutiérrez, R. A., Krouk, G., et al. (2019). Network Walking charts transcriptional dynamics of nitrogen signaling by integrating validated and predicted genome-wide interactions. Nat. Commun. 10, 1569.

Castaings, L., Camargo, A., Pocholle, D., Gaudon, V., Texier, Y., Boutet-Mercey, S., Taconnat, L., Renou, J.-P., Daniel-Vedele, F., Fernandez, E., et al. (2009). The nodule inception-like protein 7 modulates nitrate sensing and metabolism in Arabidopsis. Plant J. 57, 426–435.

Chan, E. K. F., Rowe, H. C. and Kliebenstein, D. J. (2010). Understanding the evolution of defense metabolites in Arabidopsis thaliana using genome-wide association mapping. Genetics 185, 991–1007.

Chan, E. K. F., Rowe, H. C., Corwin, J. A., Joseph, B. and Kliebenstein, D. J. (2011). Combining genome-wide association mapping and transcriptional networks to identify novel genes controlling glucosinolates in Arabidopsis thaliana. PLoS Biol. 9, e1001125.

Cheng, C.-Y., Li, Y., Varala, K., Bubert, J., Huang, J., Kim, G. J., Halim, J., Arp, J., Shih, H.-J. S., Levinson, G., et al. (2021). Evolutionarily informed machine learning enhances the power of predictive gene-to-phenotype relationships. Nat. Commun. 12, 5627.

Coleto, I., de la Peña, M., Rodríguez-Escalante, J., Bejarano, I., Glauser, G., Aparicio-Tejo, P. M., González-Moro, M. B. and Marino, D. (2017). Leaves play a central role in the adaptation of nitrogen and sulfur metabolism to ammonium nutrition in oilseed rape (Brassica napus). BMC Plant Biol. 17, 157.

Daxenbichler, M. E., Spencer, G. F., Carlson, D. G., Rose, G. B., Brinker, A. M. and Powell, R. G. (1991). Glucosinolate composition of seeds from 297 species of wild plants. Phytochemistry 30, 2623–2638.

Du, Z., Zhou, X., Ling, Y., Zhang, Z. and Su, Z. (2010). agriGO: a GO analysis toolkit for the agricultural community. Nucleic Acids Res. 38, W64–70.

Epstein, E. and Bloom, A. J. (2005). Mineral nutrition of plants: principles and perspectives. (ed. Massachusetts, S. A.).

Esteban, R., Ariz, I., Cruz, C. and Moran, J. F. (2016). Review: Mechanisms of ammonium toxicity and the quest for tolerance. Plant Sci. 248, 92–101.

Fuertes-Mendizábal, T., González-Torralba, J., Arregui, L. M., González-Murua, C., González-Moro, M. B. and Estavillo, J. M. (2013). Ammonium as sole N source improves grain quality in wheat. J. Sci. Food Agric. 93, 2162–2171.

Fujii, H. and Zhu, J.-K. (2009). Arabidopsis mutant deficient in 3 abscisic acid-activated protein kinases reveals critical roles in growth, reproduction, and stress. Proc Natl Acad Sci USA 106, 8380–8385.

Gaudinier, A., Rodriguez-Medina, J., Zhang, L., Olson, A., Liseron-Monfils, C., Bågman, A.-M., Foret, J., Abbitt, S., Tang, M., Li, B., et al. (2018). Transcriptional regulation of nitrogen-associated metabolism and growth. Nature 563, 259–264.

Gehan, M. A., Fahlgren, N., Abbasi, A., Berry, J. C., Callen, S. T., Chavez, L., Doust, A. N., Feldman, M. J., Gilbert, K. B., Hodge, J. G., et al. (2017). PlantCV v2: Image analysis software for high-throughput plant phenotyping. PeerJ 5, e4088.

Gifford, M. L., Dean, A., Gutierrez, R. A., Coruzzi, G. M. and Birnbaum, K. D. (2008). Cell-specific nitrogen responses mediate developmental plasticity. Proc Natl Acad Sci USA 105, 803–808.

Grimm, D. G., Roqueiro, D., Salomé, P. A., Kleeberger, S., Greshake, B., Zhu, W., Liu, C., Lippert, C., Stegle, O., Schölkopf, B., et al. (2017). easyGWAS: ACloud-Based Platform for Comparing the Results of Genome-Wide Association Studies. Plant Cell 29, 5–19.

Halkier, B. A. and Gershenzon, J. (2006). Biology and biochemistry of glucosinolates. Annu. Rev. Plant Biol. 57, 303–333.

Hansen, B. G., Kerwin, R. E., Ober, J. A., Lambrix, V. M., Mitchell-Olds, T., Gershenzon, J., Halkier, B. A. and Kliebenstein, D. J. (2008). Anovel 2-oxoacid-dependent dioxygenase involved in the formation of the goiterogenic 2-hydroxybut-3-enyl glucosinolate and generalist insect resistance in Arabidopsis,. Plant Physiol. 148, 2096–2108.

He, H., Liang, G., Li, Y., Wang, F. and Yu, D. (2014). Two young MicroRNAs originating from target duplication mediate nitrogen starvation adaptation via regulation of glucosinolate synthesis in Arabidopsis thaliana. Plant Physiol. 164, 853–865.

Jian, S., Liao, Q., Song, H., Liu, Q., Lepo, J. E., Guan, C., Zhang, J., Ismail, A. M. and Zhang, Z. (2018). NRT1.1-Related NH4+ Toxicity Is Associated with a Disturbed Balance between NH4+ Uptake and Assimilation. Plant Physiol. 178, 1473–1488.

Kahle, D. and Wickham, H. (2013). ggmap: Spatial Visualization with ggplot2. R J. 5, 144.

Kang, H. M., Sul, J. H., Service, S. K., Zaitlen, N. A., Kong, S.-Y., Freimer, N. B., Sabatti, C. and Eskin, E. (2010). Variance component model to account for sample structure in genome-wide association studies. Nat. Genet. 42, 348–354.

Katz, E., Nisani, S., Sela, M., Behar, H. and Chamovitz, D. A. (2015). The effect of indole-3-carbinol on PIN1 and PIN2 in Arabidopsis roots. Plant Signal. Behav. 10, e1062200.

Katz, E., Bagchi, R., Jeschke, V., Rasmussen, A. R. M., Hopper, A., Burow, M., Estelle, M. and Kliebenstein, D. J. (2020). Diverse allyl glucosinolate catabolites independently influence root growth and development. Plant Physiol. 183, 1376–1390.

Katz, E., Li, J.-J., Jaegle, B., Ashkenazy, H., Abrahams, S. R., Bagaza, C., Holden, S., Pires, C. J., Angelovici, R. and Kliebenstein, D. J. (2021). Genetic variation, environment and demography intersect to shape Arabidopsis defense metabolite variation across Europe. eLife 10,.

Kawakatsu, T., Huang, S.-S. C., Jupe, F., Sasaki, E., Schmitz, R. J., Urich, M. A., Castanon, R., Nery, J. R., Barragan, C., He, Y., et al. (2016). Epigenomic Diversity in a Global Collection of Arabidopsis thaliana Accessions. Cell 166, 492–505.

Kerwin, R., Feusier, J., Corwin, J., Rubin, M., Lin, C., Muok, A., Larson, B., Li, B., Joseph, B., Francisco, M., et al. (2015). Natural genetic variation in Arabidopsis thaliana defense metabolism genes modulates field fitness. eLife 4,.

Kliebenstein, D. J., Lambrix, V. M., Reichelt, M., Gershenzon, J. and Mitchell-Olds, T. (2001a). Gene duplication in the diversification of secondary metabolism: tandem 2-oxoglutarate-dependent dioxygenases control glucosinolate biosynthesis in Arabidopsis. Plant Cell 13, 681–693.

Kliebenstein, D. J., Kroymann, J., Brown, P., Figuth, A., Pedersen, D., Gershenzon, J. and Mitchell-Olds, T. (2001b). Genetic control of natural variation in Arabidopsis glucosinolate accumulation. Plant Physiol. 126, 811–825.

Kliebenstein, D. J., Gershenzon, J. and Mitchell-Olds, T. (2001c). Comparative quantitative trait loci mapping of aliphatic, indolic and benzylic glucosinolate production in Arabidopsis thaliana leaves and seeds. Genetics 159, 359–370.

Konishi, M. and Yanagisawa, S. (2013). Arabidopsis NIN-like transcription factors have a central role in nitrate signalling. Nat. Commun. 4, 1617.

Krouk, G., Mirowski, P., LeCun, Y., Shasha, D. E. and Coruzzi, G. M. (2010). Predictive network modeling of the high-resolution dynamic plant transcriptome in response to nitrate. Genome Biol. 11, R123.

La, G.-X. (2013). Effect of NH4+/NO3-ratios on the growth and bolting stem glucosinolate content of chinese kale (“Brassica alboglabra” L.H. bailey). Australian Journal of Crop Science 7, 618–624.

Liu, Y. and von Wirén, N. (2017). Ammonium as a signal for physiological and morphological responses in plants. J. Exp. Bot. 68, 2581–2592.

Liu, S., Zenda, T., Dong, A., Yang, Y., Wang, N. and Duan, H. (2021). Global Transcriptome and Weighted Gene Co-expression Network Analyses of Growth-Stage-Specific Drought Stress Responses in Maize. Front. Genet. 12, 645443.

Li, B., Li, G., Kronzucker, H. J., Baluška, F. and Shi, W. (2014). Ammonium stress in Arabidopsis: signaling, genetic loci, and physiological targets. Trends Plant Sci. 19, 107–114.

Malinovsky, F. G., Thomsen, M.-L. F., Nintemann, S. J., Jagd, L. M., Bourgine, B., Burow, M. and Kliebenstein, D. J. (2017). An evolutionarily young defense metabolite influences the root growth of plants via the ancient TOR signaling pathway. eLife 6,.

Marino, D. and Moran, J. F. (2019). Can ammonium stress be positive for plant performance? Front. Plant Sci. 10, 1103.

Marino, D., Ariz, I., Lasa, B., Santamaría, E., Fernández-Irigoyen, J., González-Murua, C. and Aparicio Tejo, P. M. (2016). Quantitative proteomics reveals the importance of nitrogen source to control glucosinolate metabolism in Arabidopsis thaliana and Brassica oleracea. J. Exp. Bot. 67, 3313–3323.

Matson, P. A., Naylor, R. and Ortiz-Monasterio, I. (1998). Integration of environmental, agronomic, and economic aspects of fertilizer management. Science 280, 112–115.

McCready, K., Spencer, V. and Kim, M. (2020). The importance of TOR kinase in plant development. Front. Plant Sci. 11, 16.

Krouk, G. (2015). AtNIGT1/HRS1 integrates nitrate and phosphate signals at the Arabidopsis root tip. Nat. Commun. 6, 6274.

Menz, J., Range, T., Trini, J., Ludewig, U. and Neuhäuser, B. (2018). Molecular basis of differential nitrogen use efficiencies and nitrogen source preferences in contrasting Arabidopsis accessions. Sci. Rep. 8, 3373.

Milocco, L. and Salazar-Ciudad, I. (2022). Evolution of the G Matrix under Nonlinear Genotype-Phenotype Maps. Am. Nat. 199, 420–435.

Obertello, M., Krouk, G., Katari, M. S., Runko, S. J. and Coruzzi, G. M. (2010). Modeling the global effect of the basic-leucine zipper transcription factor 1 (bZIP1) on nitrogen and light regulation in Arabidopsis. BMC Syst. Biol. 4, 111.

Omirou, M. D., Papadopoulou, K. K., Papastylianou, I., Constantinou, M., Karpouzas, D. G., Asimakopoulos, I. and Ehaliotis, C. (2009). Impact of nitrogen and sulfur fertilization on the composition of glucosinolates in relation to sulfur assimilation in different plant organs of broccoli. J. Agric. Food Chem. 57, 9408–9417.

Pigliucci, M. and Murren, C. J. (2003). Perspective: Genetic assimilation and a possible evolutionary paradox: can macroevolution sometimes be so fast as to pass us by? Evolution 57, 1455–1464.

Pigliucci, M., Murren, C. J. and Schlichting, C. D. (2006). Phenotypic plasticity and evolution by genetic assimilation. J. Exp. Biol. 209, 2362–2367.

Pound, M. P., French, A. P., Atkinson, J. A., Wells, D. M., Bennett, M. J. and Pridmore, T. (2013). RootNav: navigating images of complex root architectures. Plant Physiol. 162, 1802–1814.

Rodman, J. E. (1980). Population variation and hybridization in sea-rockets (cakile, cruciferae): seed glucosinolate characters. Am. J. Bot. 67, 1145–1159.

Rosas, U., Cibrian-Jaramillo, A., Ristova, D., Banta, J. A., Gifford, M. L., Fan, A. H., Zhou, R. W., Kim, G. J., Krouk, G., Birnbaum, K. D., et al. (2013). Integration of responses within and across Arabidopsis natural accessions uncovers loci controlling root systems architecture. Proc Natl Acad Sci USA 110, 15133–15138.

Rubin, G., Tohge, T., Matsuda, F., Saito, K. and Scheible, W.-R. (2009). Members of the LBD family of transcription factors repress anthocyanin synthesis and affect additional nitrogen responses in Arabidopsis. Plant Cell 21, 3567–3584.

Salehin, M., Li, B., Tang, M., Katz, E., Song, L., Ecker, J. R., Kliebenstein, D. J. and Estelle, M. (2019). Auxin-sensitive Aux/IAA proteins mediate drought tolerance in Arabidopsis by regulating glucosinolate levels. Nat. Commun. 10, 4021.

Schlichting, C. D. and Pigliucci, M. (1993). Control of phenotypic plasticity via regulatory genes. Am. Nat. 142, 366–370.

Schraiber, J. G. and Landis, M. J. (2015). Sensitivity of quantitative traits to mutational effects and number of loci. Theor. Popul. Biol. 102, 85–93.

Slaten, M. L., Yobi, A., Bagaza, C., Chan, Y. O., Shrestha, V., Holden, S., Katz, E., Kanstrup, C., Lipka, A. E., Kliebenstein, D. J., et al. (2020). mGWAS Uncovers Gln-Glucosinolate Seed-Specific Interaction and its Role in Metabolic Homeostasis. Plant Physiol. 183, 483–500.

Sønderby, I. E., Geu-Flores, F. and Halkier, B. A. (2010). Biosynthesis of glucosinolates--gene discovery and beyond. Trends Plant Sci. 15, 283–290.

The 1001 Genomes Consortium (2016). 1,135 Genomes reveal the global pattern of polymorphism in Arabidopsis thaliana. Cell 166, 481–491.

Tian, Q.-Y., Sun, P. and Zhang, W.-H. (2009). Ethylene is involved in nitrate-dependent root growth and branching in Arabidopsisthaliana. New Phytol. 184, 918–931.

Turner, S. D. (2014). qqman: an R package for visualizing GWAS results using Q-Q and manhattan plots. BioRxiv.

Urbut, S. M., Wang, G., Carbonetto, P. and Stephens, M. (2019). Flexible statistical methods for estimating and testing effects in genomic studies with multiple conditions. Nat. Genet. 51, 187–195.

Varala, K., Marshall-Colón, A., Cirrone, J., Brooks, M. D., Pasquino, A. V., Léran, S., Mittal, S., Rock, T. M., Edwards, M. B., Kim, G. J., et al. (2018). Temporal transcriptional logic of dynamic regulatory networks underlying nitrogen signaling and use in plants. Proc Natl Acad Sci USA 115, 6494–6499.

Vidal, E. A., Álvarez, J. M. and Gutiérrez, R. A. (2014). Nitrate regulation ofAFB3 andNAC4 gene expression inArabidopsis roots depends on NRT1.1 nitrate transport function. Plant Signal. Behav. 9, e28501.

Waddington, C. H. (1953). Genetic assimilation of an acquired character. Evolution 7, 118–126.

Waidmann, S., Sarkel, E. and Kleine-Vehn, J. (2020). Same same, but different: growth responses of primary and lateral roots. J. Exp. Bot. 71, 2397–2411.

Wickham, H., Averick, M., Bryan, J., Chang, W., McGowan, L., François, R., Grolemund, G., Hayes, A., Henry, L., Hester, J., et al. (2019). Welcome to the tidyverse. JOSS 4, 1686.

Wisecaver, J. H., Borowsky, A. T., Tzin, V., Jander, G., Kliebenstein, D. J. and Rokas, A. (2017). Aglobal coexpression network approach for connecting genes to specialized metabolic pathways in plants. Plant Cell 29, 944–959.

Wright, S. I., Lauga, B. and Charlesworth, D. (2002). Rates and patterns of molecular evolution in inbred and outbred Arabidopsis. Mol. Biol. Evol. 19, 1407–1420.

Xu, N., Wang, R., Zhao, L., Zhang, C., Li, Z., Lei, Z., Liu, F., Guan, P., Chu, Z., Crawford, N. M., et al. (2016). The Arabidopsis NRG2 Protein Mediates Nitrate Signaling and Interacts with and Regulates Key Nitrate Regulators. Plant Cell 28, 485–504.

Yan, X. and Chen, S. (2007). Regulation of plant glucosinolate metabolism. Planta 226, 1343–1352.

Zhang, H. and Forde, B. G. (1998). An Arabidopsis MADS box gene that controls nutrient-induced changes in root architecture. Science 279, 407–409.

Zhang, X., Davidson, E. A., Mauzerall, D. L., Searchinger, T. D., Dumas, P. and Shen, Y. (2015). Managing nitrogen for sustainable development. Nature 528, 51–59.

