## Supplemental figures for "Differential genetic variation underlying Ammonium and Nitrate responses in *Arabidopsis thaliana*"

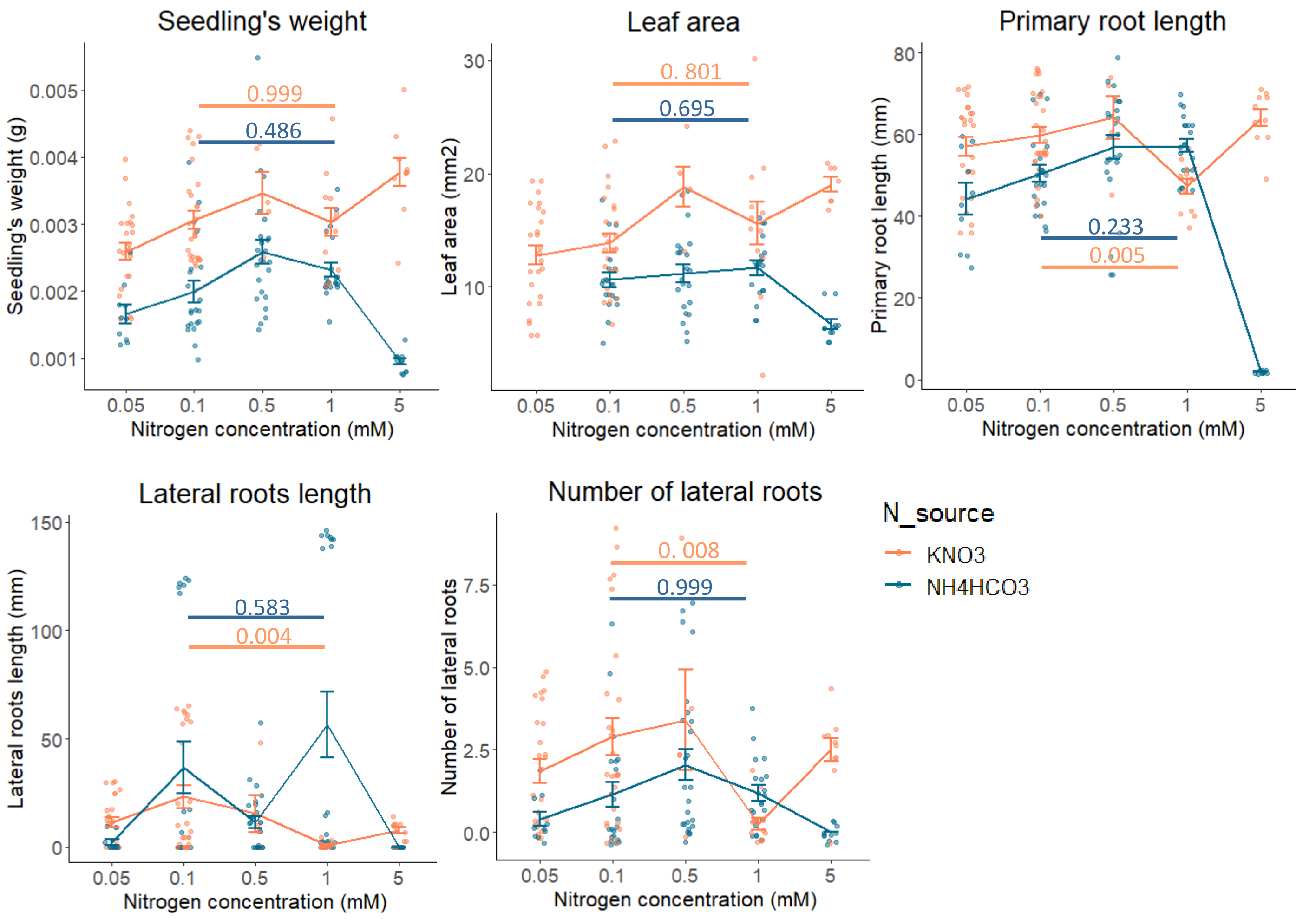


**Supp figure 1: Col-0 seedlings present different phenotypes when grown on different nitrogen conditions:** Col-0 seedlings were grown on MS medium with nitrate (KNO3) or ammonium (NH4HCO3) as a sole nitrogen source in different concentrations (0.05-5mM). After 12 days different developmental phenotypes were measured. The indicated values indicate differences between phenotype values of seedlings grown on 0.1mM and 1mM nitrate (orange) or ammonium (blue), that were tested via ANOVA followed by Tukey's honestly significant (HSD) mean-separation test for each nitrogen source separately. Error bars represent means ± standard errors. The jitter function used for these plots adds a small amount of random variation to the location of each point to ovoid overplotting.

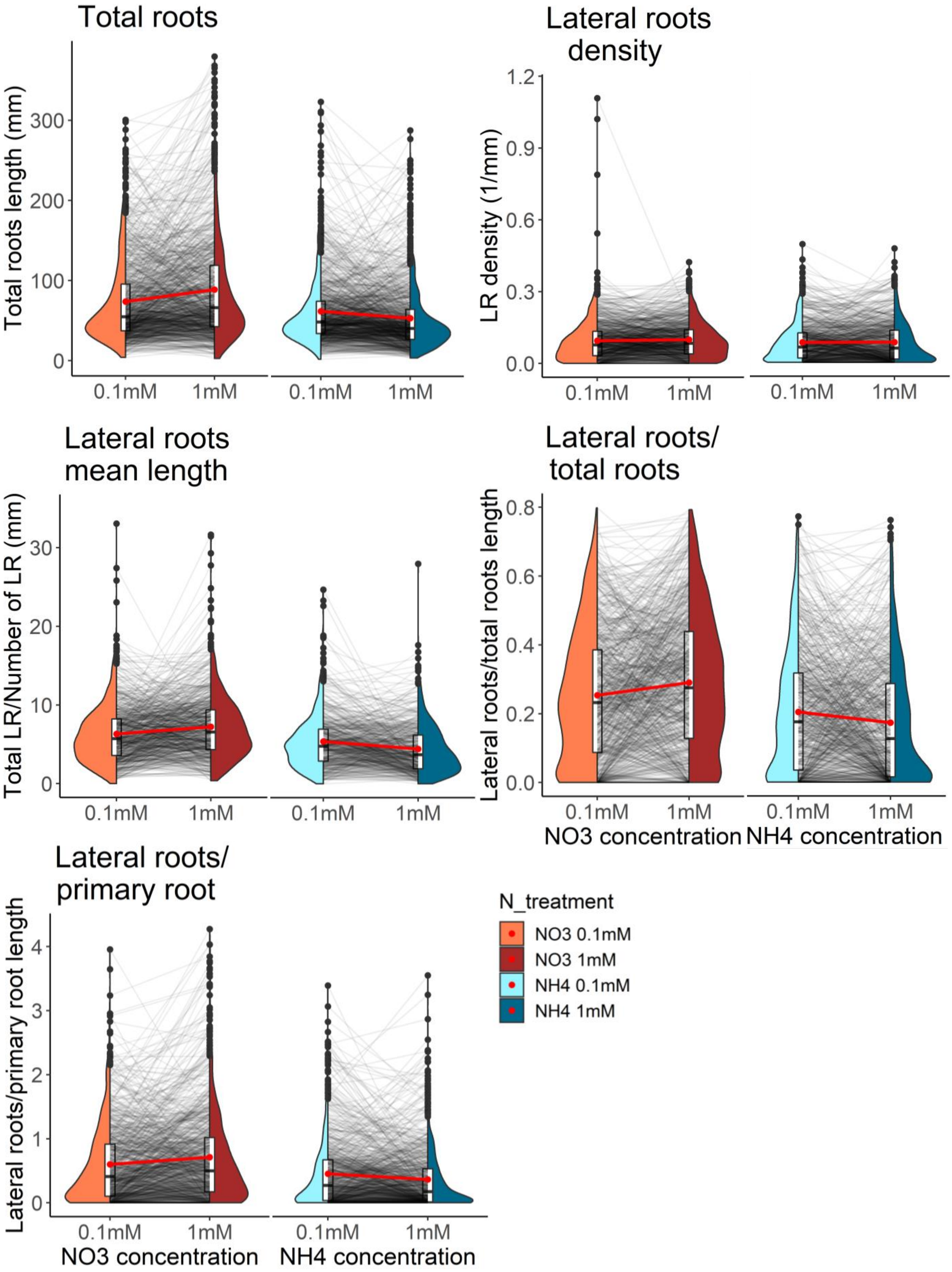


**Supp figure 2: Nitrogen conditions affect different traits across natural accessions:** phenotypes of Arabidopsis accessions grown on 4 different nitrogen conditions (nitrate 0.1mM, nitrate 1mM, ammonium 0.1mM, ammonium 1mM). Grey lines between each pair of nitrogen concentrations connect the means of each individual accession grown under each nitrogen concentration. Red lines between the violins connect the mean phenotype between two concentrations of the same nitrogen source. Lateral roots density= (Number of lateral roots/Primary root length).

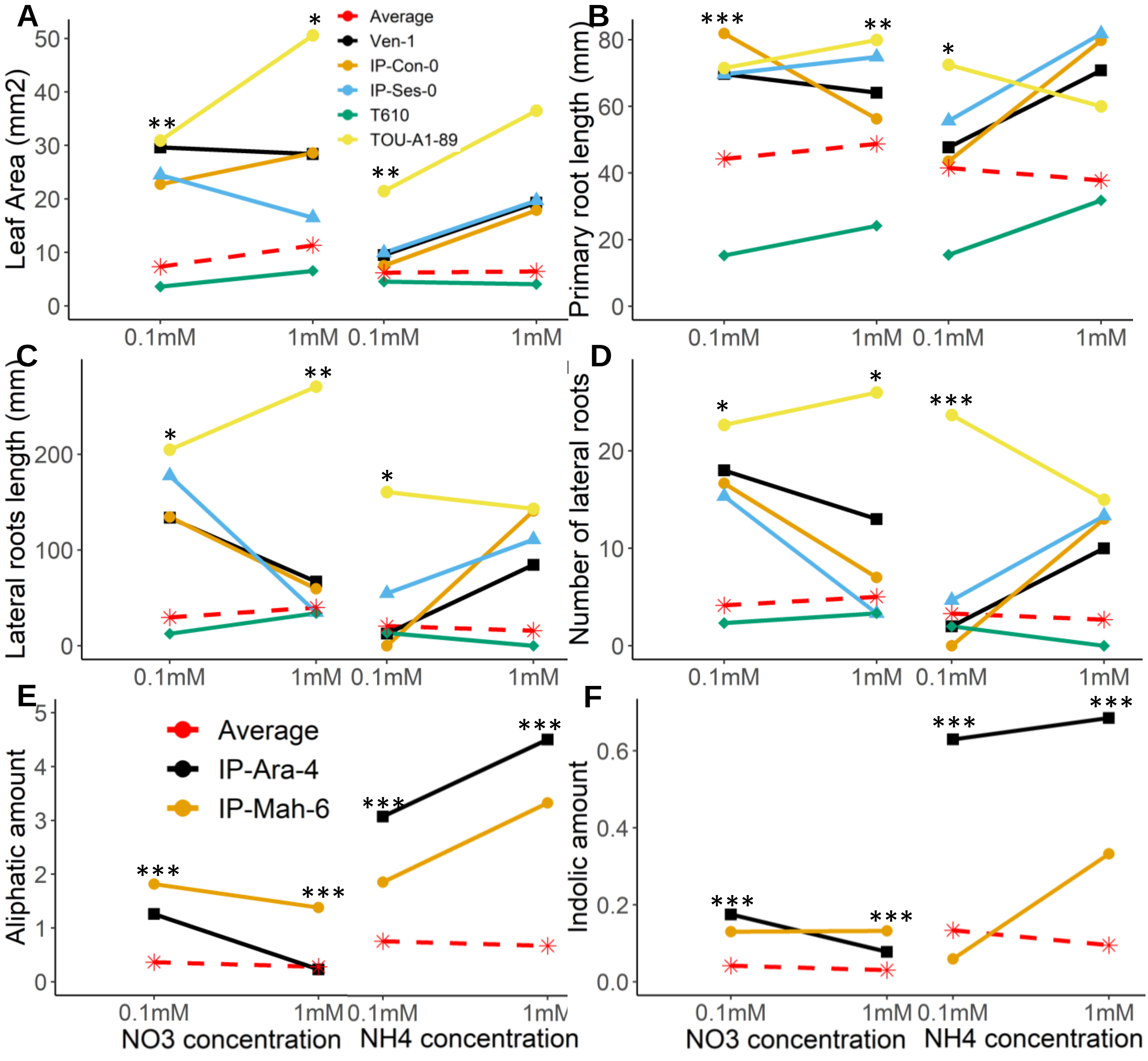


**Supp figure 3: Illustrative Arabidopsis accessions show diverse phenotypes under different nitrogen conditions:** phenotypes of Arabidopsis accessions grown on 4 different nitrogen conditions (nitrate 0.1mM, nitrate 1mM, ammonium 0.1mM, ammonium 1mM). A. Leaf area. B. Primary root length. C. Lateral roots length. D. Number of lateral roots. E. Aliphatic GSLs. F. Indolic GSLs. Aliphatic and indolic GSLs amounts were normalized by total roots length for seedlings grown on NO3 1mM and NH4 1mM (amount units are umol/mm), and by total roots length and leaves area for seedlings grown on NO3 0.1mM and NH4 0.1mM (amount units are umol/mm+mm2). Significance was tested via two-way ANOVAs that was done for each measured trait under each nitrogen condition against the accession (*$P < 0.05$, **$P < 0.01$, and ***$P < 0.0001$). Detailed statistic in Table S9.

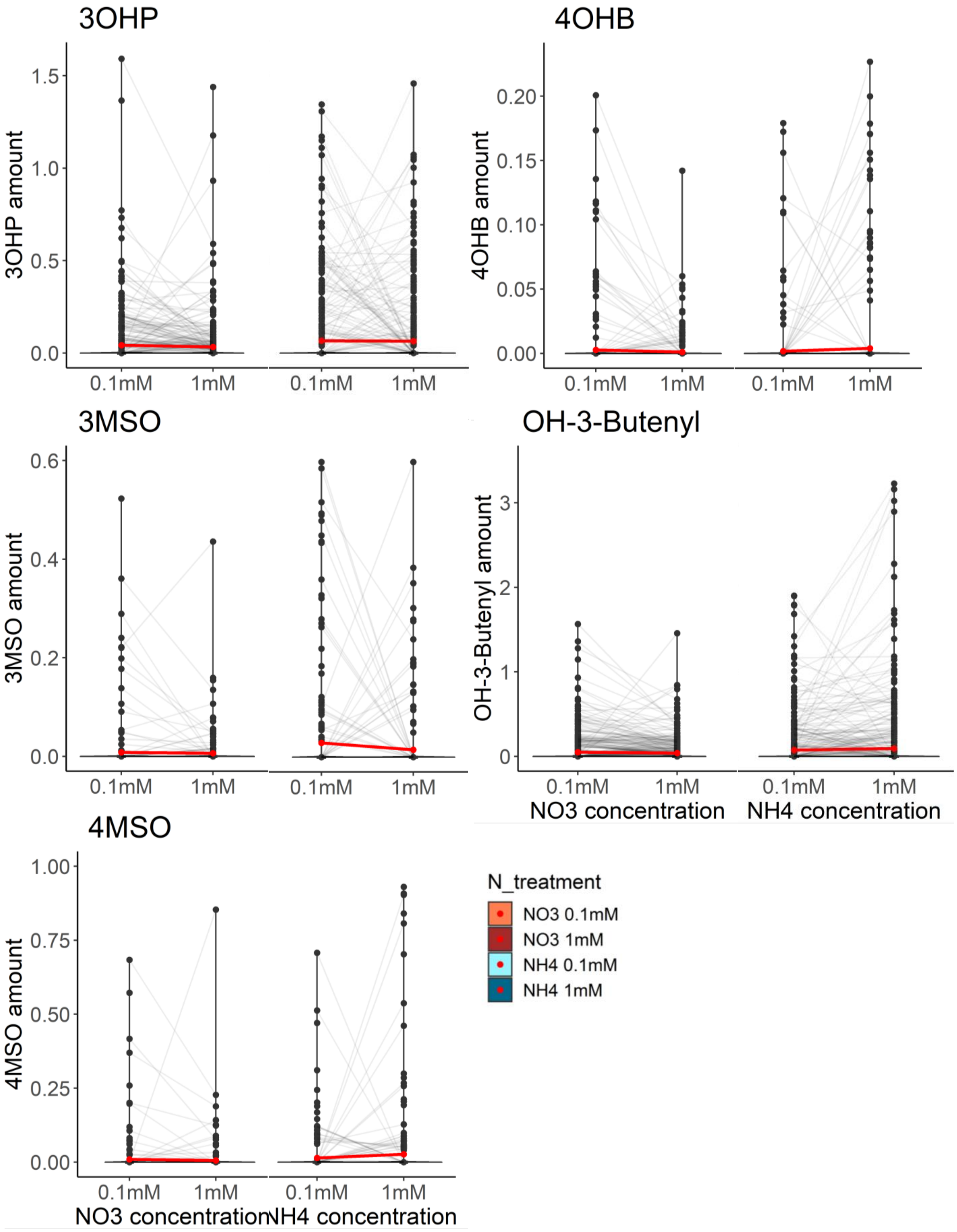


**Supp figure 4: Nitrogen conditions affect different traits across natural accessions:** phenotypes of Arabidopsis accessions grown on 4 different nitrogen conditions (nitrate 0.1mM, nitrate 1mM, ammonium 0.1mM, ammonium 1mM). Grey lines between each pair of nitrogen concentrations connect the means of each individual accession grown under each nitrogen concentration. Red lines between the violins connect the mean phenotype between two concentrations of the same nitrogen source. Aliphatic and indolic GSLs amounts were normalized by total roots length for seedlings grown on NO3 1mM and NH4 1mM (amount units are umol/mm), and by total roots length and leaves area for seedlings grown on NO3 0.1mM and NH4 0.1mM (amount units are umol/mm+mm2).

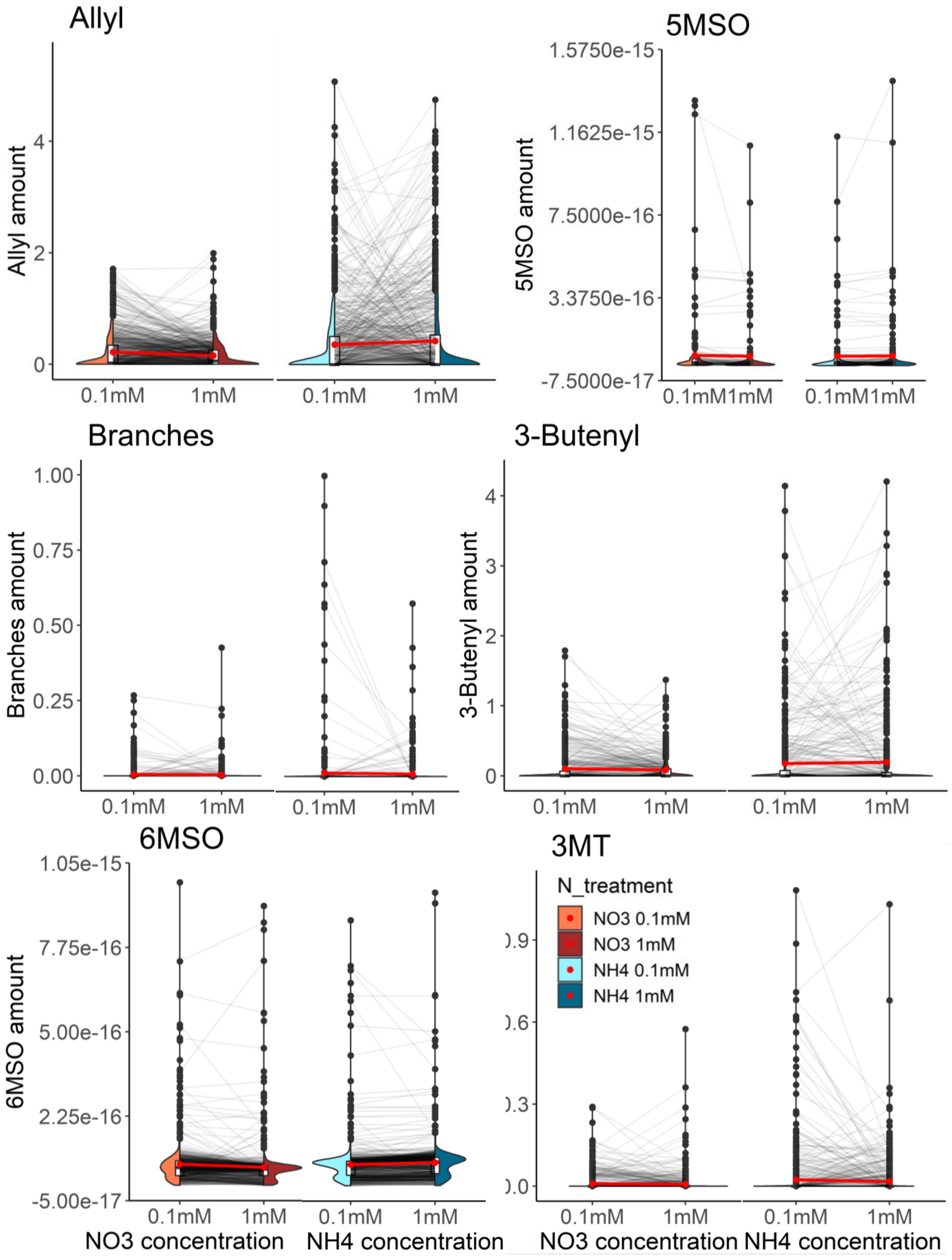


**Supp figure 5: Nitrogen conditions affect different traits across natural accessions:** phenotypes of Arabidopsis accessions grown on 4 different nitrogen conditions (nitrate 0.1mM, nitrate 1mM, ammonium 0.1mM, ammonium 1mM). Grey lines between each pair of nitrogen concentrations connect the means of each individual accession grown under each nitrogen concentration. Red lines between the violins connect the mean phenotype between two concentrations of the same nitrogen source. Aliphatic and indolic GSLs amounts were normalized by total roots length for seedlings grown on NO3 1mM and NH4 1mM (amount units are umol/mm), and by total roots length and leaves area for seedlings grown on NO3 0.1mM and NH4 0.1mM (amount units are umol/mm+mm2).

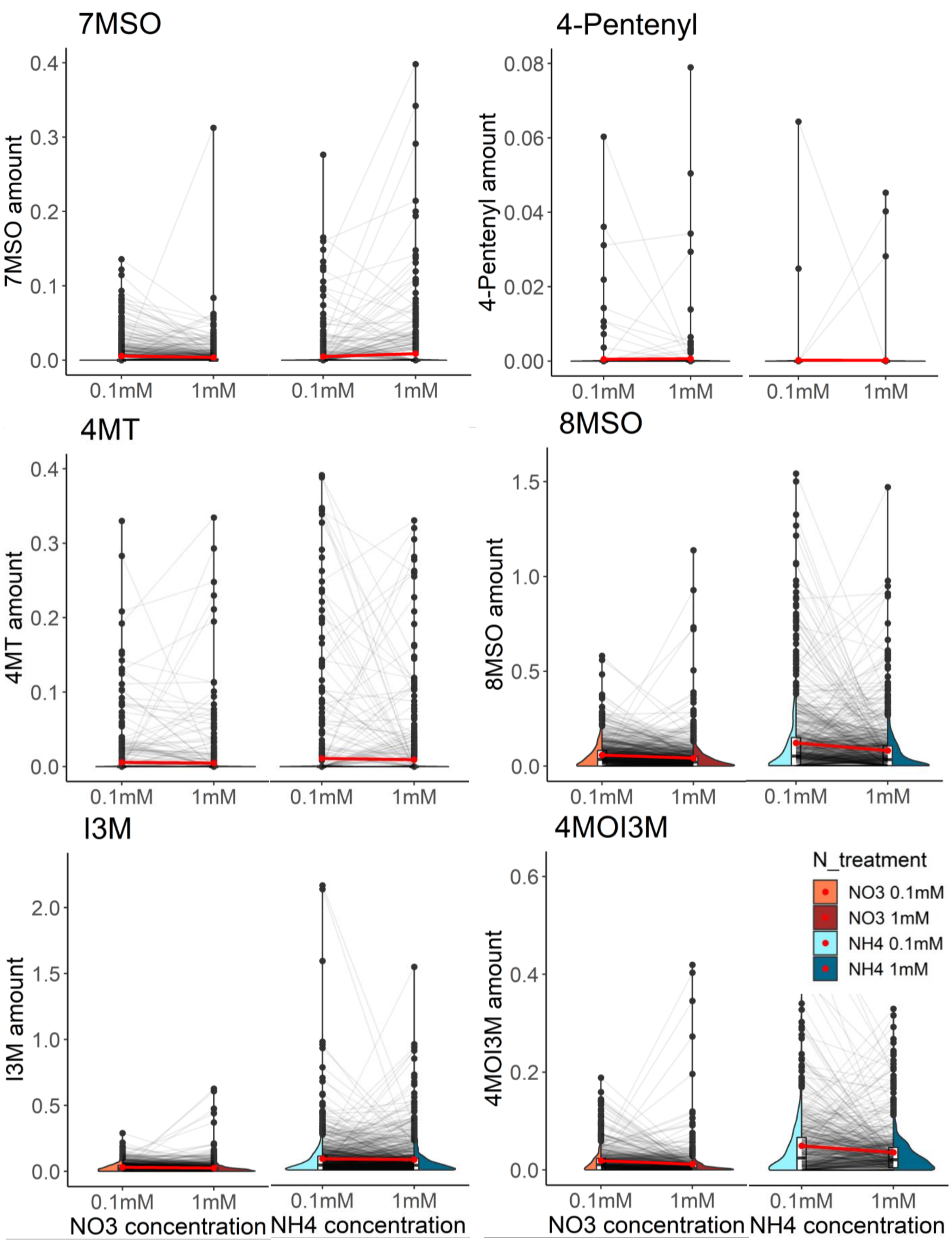


**Supp figure 6: Nitrogen conditions affect different traits across natural accessions:** phenotypes of Arabidopsis accessions grown on 4 different nitrogen conditions (nitrate 0.1mM, nitrate 1mM, ammonium 0.1mM, ammonium 1mM). Grey lines between each pair of nitrogen concentrations connect the means of each individual accession grown under each nitrogen concentration. Red lines between the violins connect the mean phenotype between two concentrations of the same nitrogen source. Aliphatic and indolic GSLs amounts were normalized by total roots length for seedlings grown on NO3 1mM and NH4 1mM (amount units are umol/mm), and by total roots length and leaves area for seedlings grown on NO3 0.1mM and NH4 0.1mM (amount units are umol/mm+mm2).

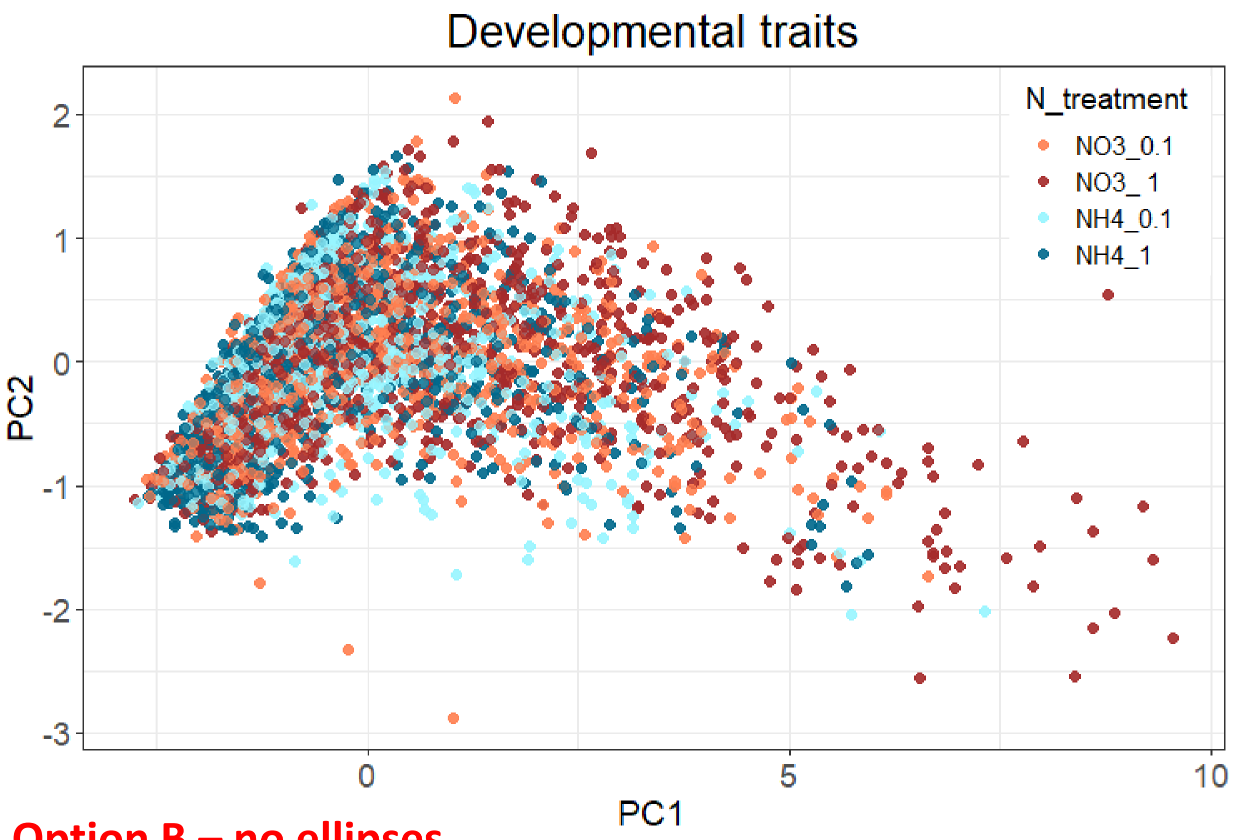


**Option B – no ellipses**

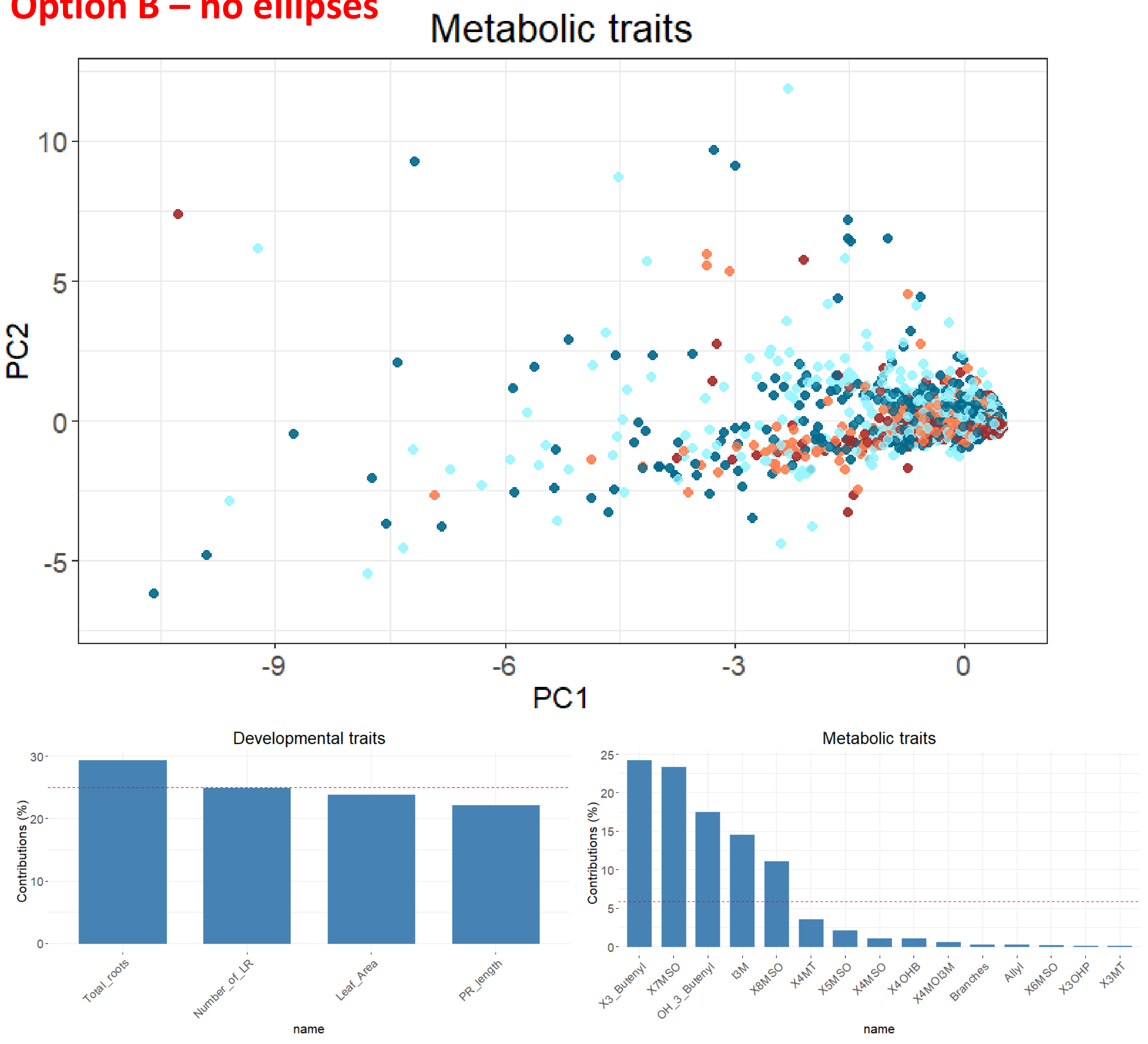


**Supp figure 7: Principal component analyses:** PCA was performed using the developmental traits (A) and individual GSLs as metabolic traits (B). C. Contribution of the individual traits to PC1 for each analysis.

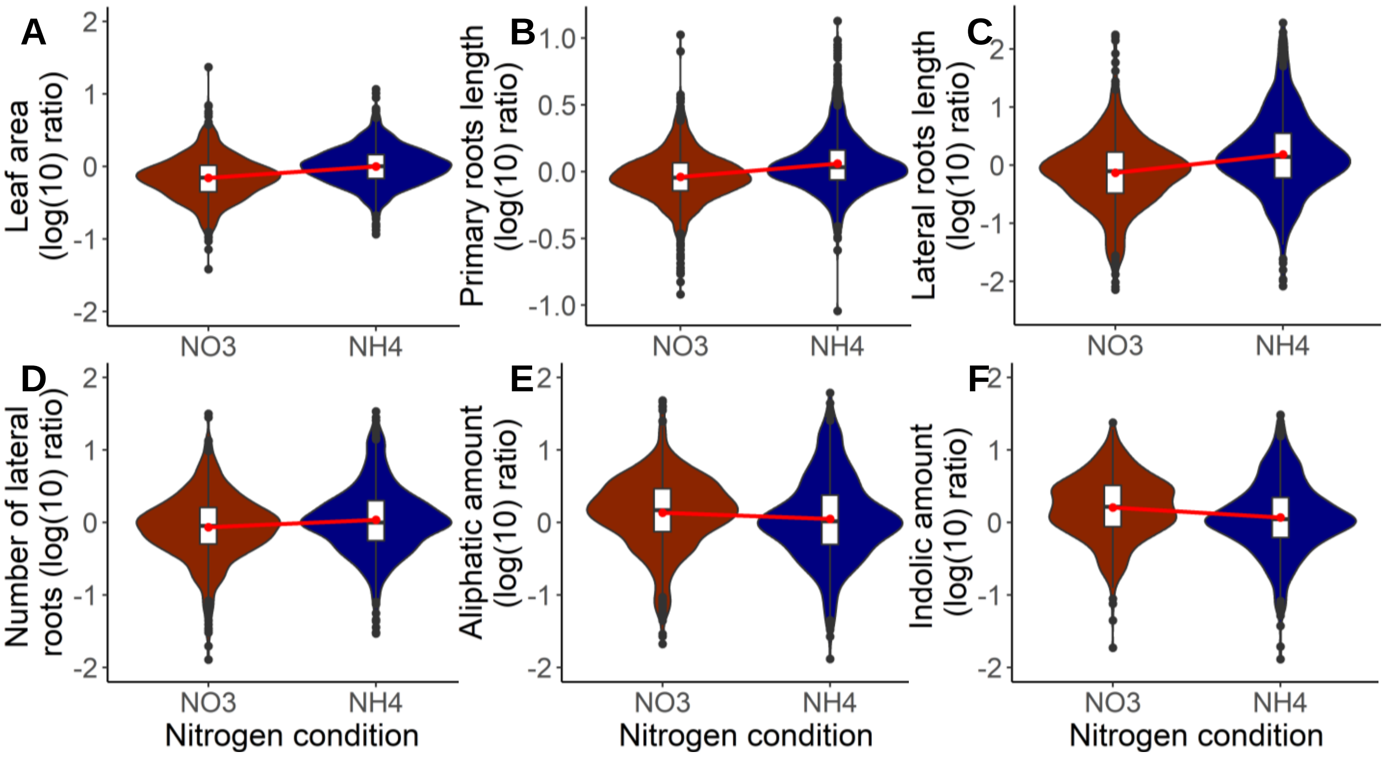


**Supp figure 8: Nitrogen conditions affect different traits across natural accessions:** the ratio between phenotypes of two concentrations of the same nitrogen source were calculated, logged and plotted for Arabidopsis accessions. The ratio was calculated between 0.1mM to 1mM for each source (nitrate and ammonium), then those values were logged. A. Leaf area. B. Primary root length. C. Lateral roots length. D. Number of lateral roots. E. Aliphatic GSLs. F. Indolic GSLs. Red lines between the violins connect the mean phenotype between the two nitrogen sources. Aliphatic and indolic GSLs amounts were normalized by total roots length for seedlings grown on NO3 1mM and NH4 1mM (amount units are umol/mm), and by total roots length and leaves area for seedlings grown on NO3 0.1mM and NH4 0.1mM (amount units are umol/mm+mm2).

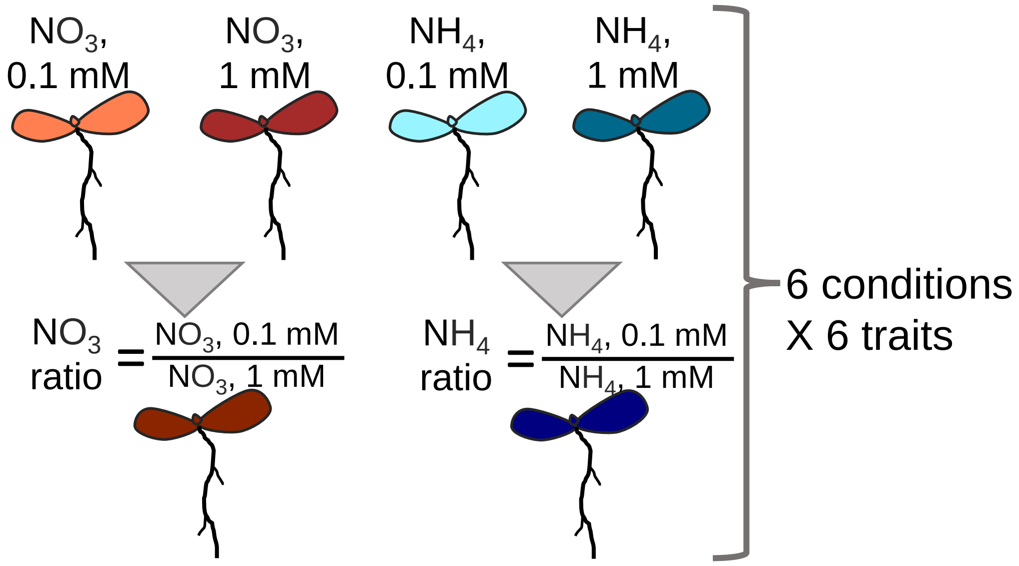


**Supp figure 9: Traits used for GWAS:** Four developmental traits and two metabolic traits (aliphatic and indolic) under each of the nitrogen conditions were used for GWAS: $NO_3$ 0.1 mM, $NO_3$ 1 mM and the logged ratio between them, $NH_4$ 0.1 mM, $NH_4$ 1 mM and the logged ratio between them. This yield 36 different GWAS (six traits, four nitrogen conditions, two ratios).

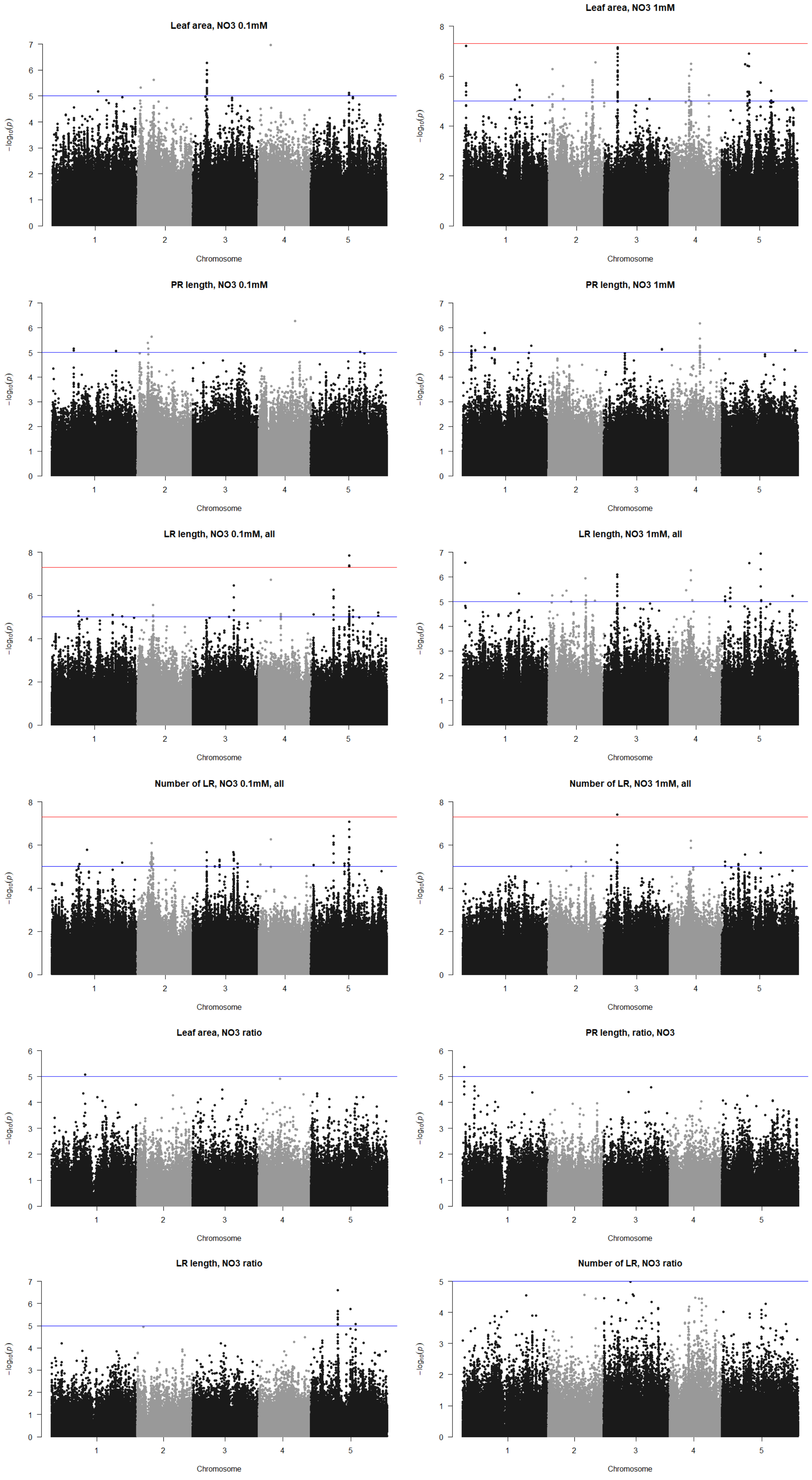


**Supp figure 10: Manhattan plots of genome-wide association analyses of developmental traits of plants grown under NO3 concentrations:** Horizontal lines represent 5% significance thresholds using Bonferroni (red) and Benjamini–Hochberg (blue).

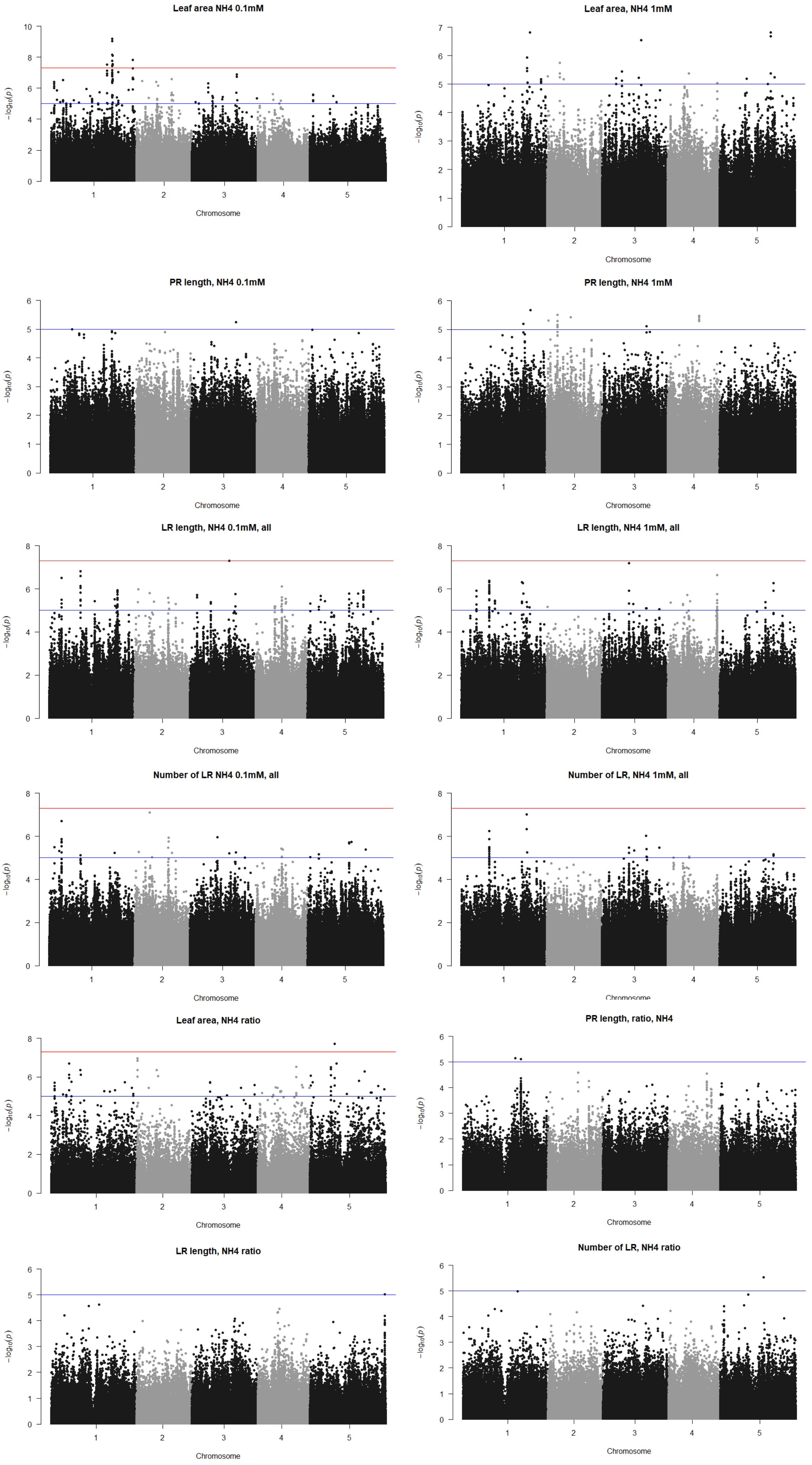


**Supp figure 11: Manhattan plots of genome-wide association analyses of developmental traits of plants grown under NH4 concentrations**: Horizontal lines represent 5% significance thresholds using Bonferroni (red) and Benjamini–Hochberg (blue).

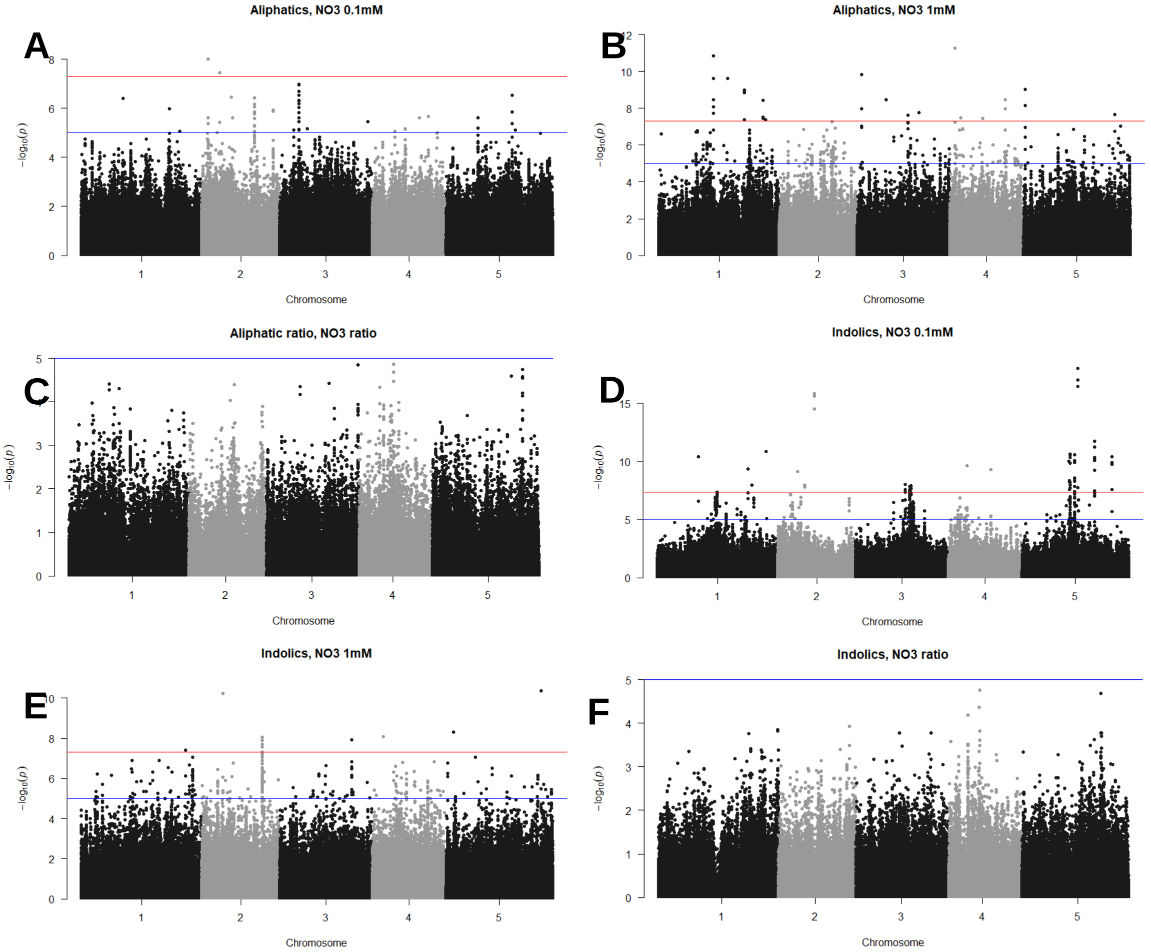


**Supp figure 12: Manhattan plots of genome-wide association analyses of aliphatic and indolic traits of plants grown under NO3 concentrations**: A. Aliphatic GSLs, NO3 0.1mM. B. Aliphatic GSLs, NO3 1mM. C. Aliphatic GSLs, NO3 ratio. D. Indolic GSLs, NO3 0.1mM. E. Indolic GSLs, NO3 1mM. F. Indolic GSLs, NO3 ratio. Horizontal lines represent 5% significance thresholds using Bonferroni (red) and Benjamini–Hochberg (blue).

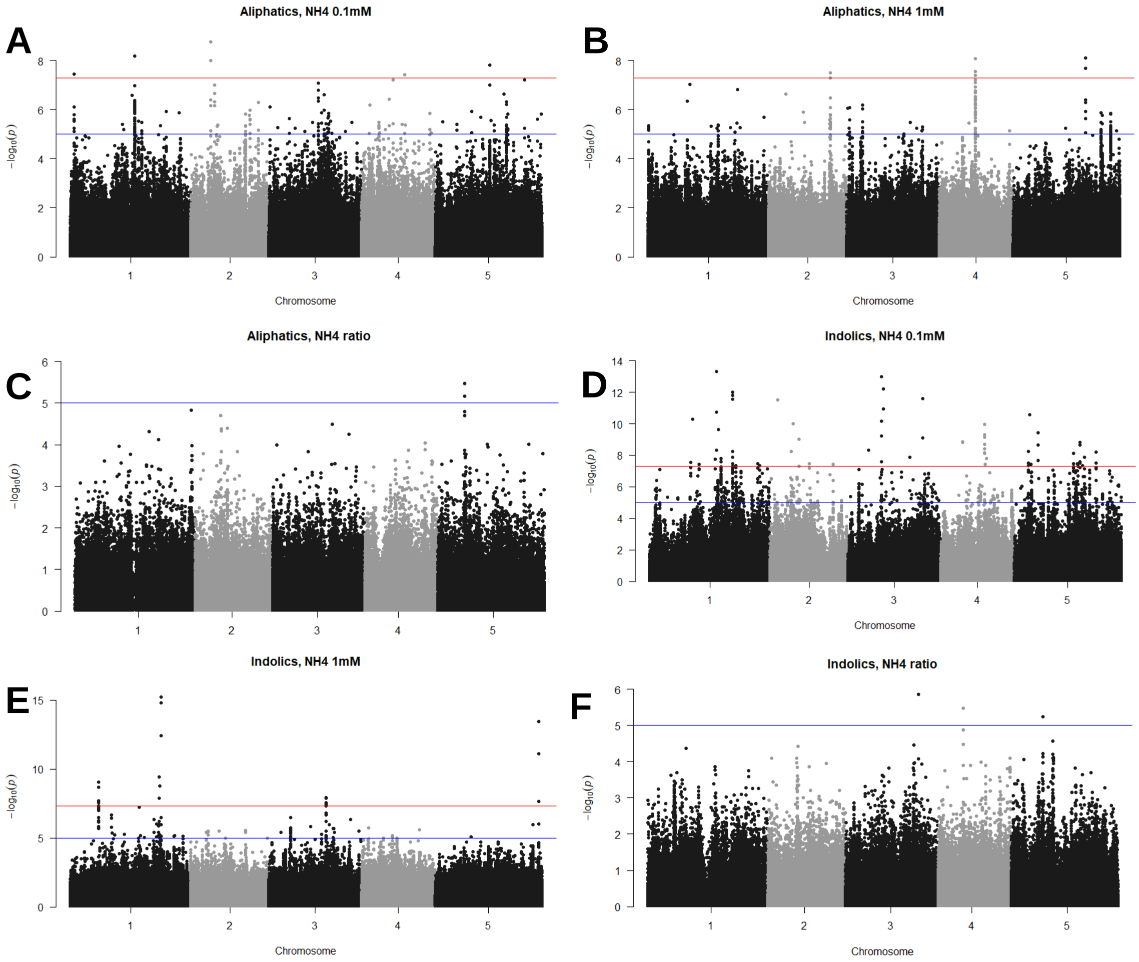


**Supp figure 13: Manhattan plots of genome-wide association analyses of aliphatic and indolic traits of plants grown under NH4 concentrations:** A. Aliphatic GSLs, NH4 0.1mM. B. Aliphatic GSLs, NH4 1mM. C. Aliphatic GSLs, NH4 ratio. D. Indolic GSLs, NH4 0.1mM. E. Indolic GSLs, NH4 1mM. F. Indolic GSLs, NH4 ratio. Horizontal lines represent 5% significance thresholds using Bonferroni (red) and Benjamini–Hochberg (blue).

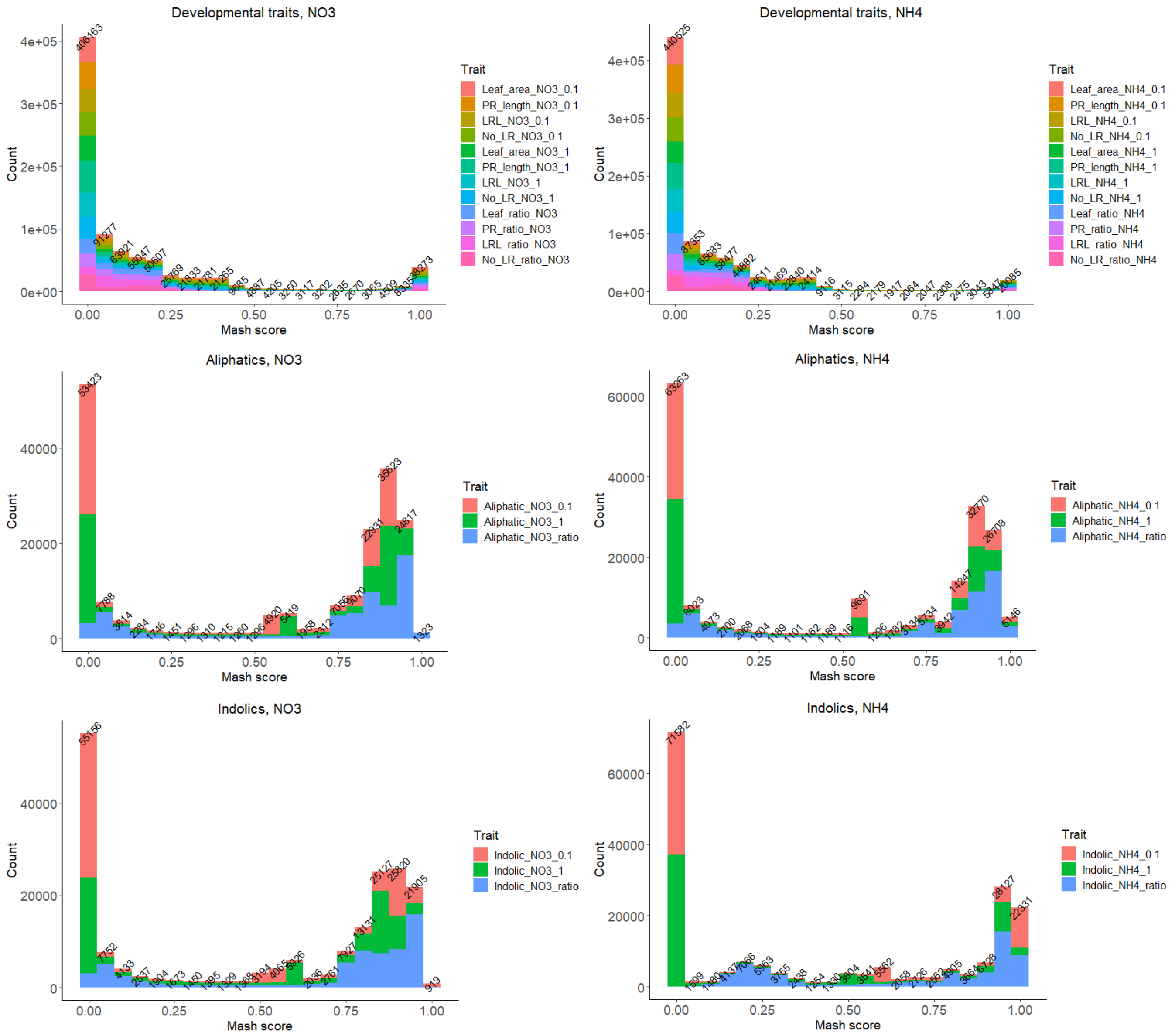


**Supp figure 14: Distribution of number of SNPs by Mash scores:** Mash analyses was done for each dataset as indicated, and the mash scores attributed from the different GWAS results are indicated. SNPs with mash score > 0.98 were chosen for further analyses.

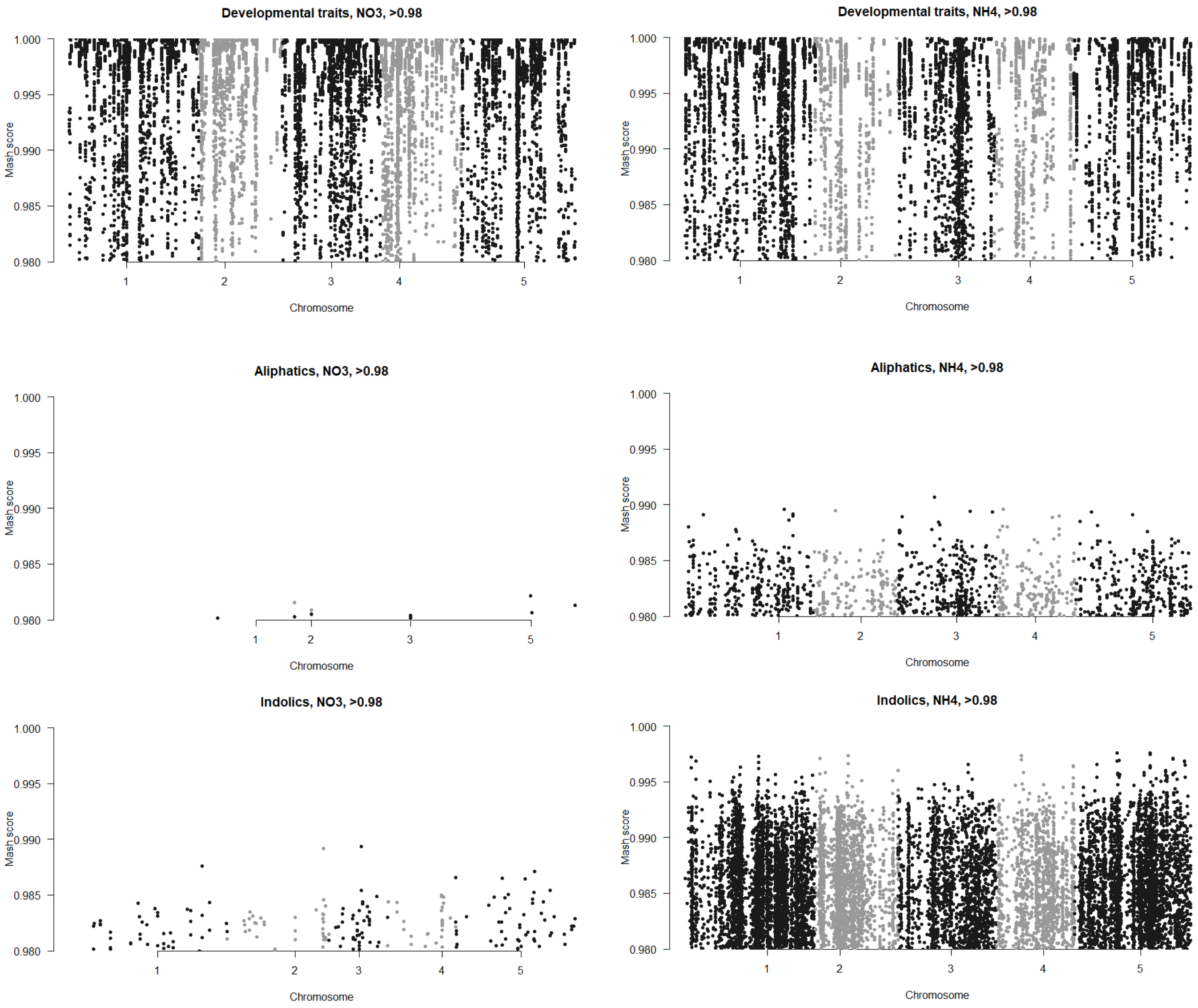


**Supp figure 15: Loci associated with nitrogen response:** SNPs yield form the MASH analyses plotted by their chromosomal position.

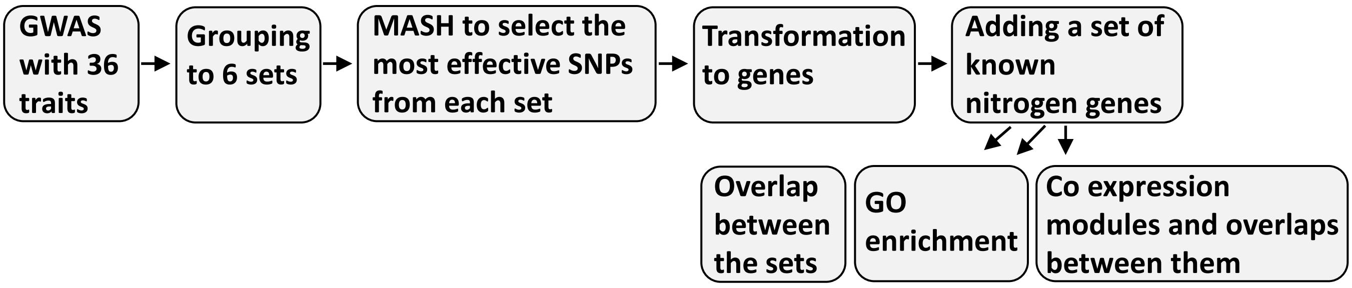


**Supp figure 16: Flow chart of the steps were taken to detect and analyze genes associated with nitrogen responses.**

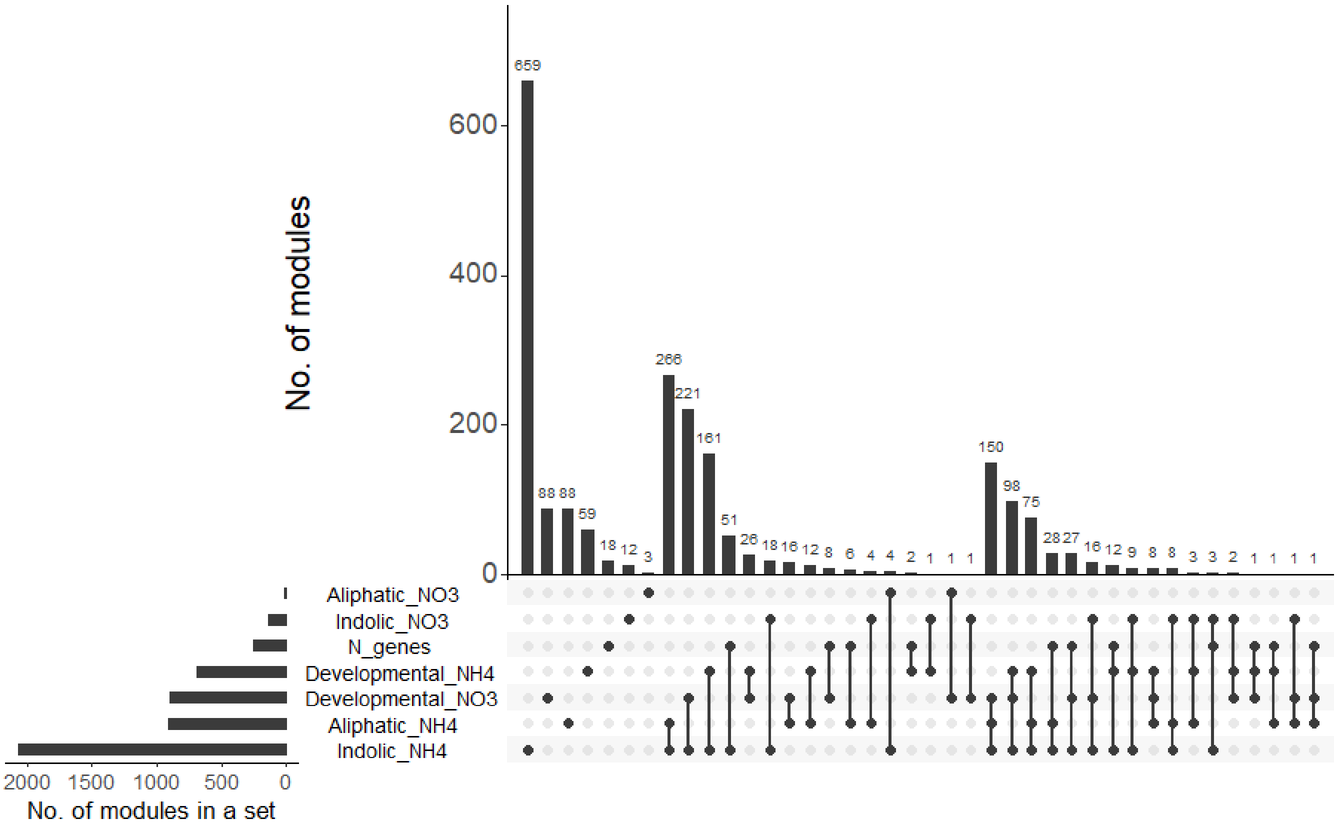


**Supp figure 17: Overlaps between gene modules yield from the different mash sets:** Genes from each set were assigned to modules, and the overlap of these modules between the different sets was analyzed. N genes are a set of Arabidopsis genes known to be related to some nitrogen process. Fisher's exact test was performed for each pair of sets (supp. Table 12). Total number of modules = 2864.

| Traits and conditions | % of overlapping genes | % of overlapping genes, including N genes | % of overlapping modules | % of overlapping modules, including N modules |
|---|---|---|---|---|
| Developmental $NO_3$ | 32.8 | 33 | 89.3 | 90 |
| Developmental $NH_4$ | 35.5 | 36 | 81.2 | 92 |
| Aliphatic $NO_3$ | 64 | 64 | 85 | 85 |
| Aliphatic $NH_4$ | 33 | 34 | 89.5 | 90 |
| Indolic $NO_3$ | 32.5 | 32.5 | 92 | 92 |
| Indolic $NH_4$ | 15 | 15.5 | 65.5 | 68 |

**Supp table 11: Percentage of overlapping genes/ modules in the different sets:** the percentage of overlapping genes/ modules was calculated with and without the list of nitrogen genes/ modules.

**List of supplemental tables**

**Supp table 1: Calibration's data**: Col-0 data that used for calibration nitrogen concentrations.
**Supp table 2: Calibration's ANOVAs**: linear models followed by two-way ANOVAs were done for each measured trait for the Col-0 data.
**Supp table 3**: **Raw data**: accessions, developmental and glucosinolate (GSL) data.
**Supp table 4**: **Emmeans data**: accessions, developmental and glucosinolate (GSL) mean data.
**Supp table 5: list of GSLs and structures.**
**Supp table 6**: **Heritability values**: percentage of variance explained by each variable was estimated for each trait based on a linear model followed by ANOVA with the indicated effects upon the measured trait.
**Supp table 7**: **P values of the different variables**: linear model with the indicated variables followed by ANOVA was conducted for each trait. The P-values were summarized in the table.
**Supp table 8**: **Differences between pairs of treatments**: differences between the indicated pairs of treatments were analyzed for each trait using aTukeyHSD test.
**Supp table 9**: **Samples' statistics**: linear models followed by two-way ANOVAs were done for each measured trait under each nitrogen condition for the selected accessions, followed by Tukey test with 95% confidence level.
**Supp table 10**: **Lists of genes yield from the different mash sets**: SNPs were transformed to genes considering 2K bps before and after the start and end point of each gene.
**Supp table 11: Percentage of overlapping genes/ modules in the different sets:** the percentage of overlapping genes/ modules was calculated with and without the list of nitrogen genes/ modules.
**Supp table 12**: **Fisher's exact test for each pair of sets**. Total number of genes = 28496.
**Supp table 13**: **List of N genes.**
**Supp table 14: Combined gene list and GO enrichment.** Columns C-H: number of SNPs in each gene under each set. Columns J-K: total number of SNPs in each gene with (K) or without (J) the N genes. Column L: number of sets each gene appeared in.
**Supp table 15: GO enrichment for the six sets.** The analyses were done using the tair tool (column A) or Agrigo tool (column D).
**Supp table 16: Lists of genes in common modules**: list of modules that are common across at least 5 sets, and the genes in those modules.
**Supp table 17: enriched GO categories using genes from the common modules.** The analyses were done using the tair tool.
